# The heritability of reinforcement learning parameters and their association with anxiety

**DOI:** 10.64898/2026.05.25.727613

**Authors:** Tim Kerr, Kirstin L. Purves, Thomas McGregor, Tom J. Barry, Kathryn J. Lester, Oliver J. Robinson, Thalia C. Eley

## Abstract

Impaired learning that both novel and previously dangerous stimuli are safe (safety and extinction learning, respectively) are long standing, robust, and heritable features of anxiety disorders, representing potential endophenotypes. The computational mechanisms underpinning them have demonstrated associations with anxiety severity in recent studies. We undertook a pre-registered replication in a tenfold larger independent sample of twins (*n* = 925). Extinction learning rates were associated with anxiety severity (*ρ*_replication_ = −0.14, *BF*_*r*0_ = 1189. 67) but safety learning rates were not. Conversely, although safety learning rates showed modest heritability (*h*^*2*^_*safety*_ = 0.16), extinction learning rates were not heritable. Accordingly, we were unable to identify genetic overlap between anxiety and either learning rate. Although this suggests neither learning rate is an anxiety endophenotype, we confirmed a cognitive-behavioral mechanism underpinning a robust marker of anxiety severity. Furthermore, we demonstrated heritability of a computationally modelled learning parameter, a key step towards establishing its biological basis.

## Introduction

Anxiety disorders are debilitating, common, chronic conditions which onset early (Bandelow & Michaelis, 2015; Bystritsky, 2006). Despite their substantial and rising prevalence (Chen et al., 2025; Morris et al., 2025), and considerable advances in the genetics of anxiety(Skelton et al., 2025; Strom et al., 2026), the cognitive-behavioural mechanisms that might mediate genetic influences on anxiety are yet to be understood (Chen et al., 2025; Morris et al., 2025). The lack of studies exploring genetic influences on relevant cognitive-behavioural processes provide limited potential endophenotypes to test in genomic studies (Penninx et al., 2021).

Fear conditioning paradigms directly quantify differences in threat responding, long understood to be instrumental in anxiety disorder development, corroborated by consistent demonstrations of associations with anxiety severity (Beckers et al., 2023; Duits et al., 2015; Kausche et al., 2025; Lissek et al., 2005; McGregor et al., 2021). These paradigms begin with an acquisition phase, where a neutral cue is paired with contemporaneous aversive experiences, counterbalanced with an unpaired control cue. Participants’ physiological or subjective fear responding to the presentation of either cue can be measured through a selection of instrumental modalities - typically electrodermal, pupillometry, cardiopulmonary, or self-report, before descriptive outcome measures of responding towards either cue are calculated. In an acquisition phase, those with anxiety disorders demonstrate increased responding (i.e., fear) to the “safe” control cue due to impaired safety learning, but no differences in responding to the “threat” cue (Kausche et al., 2025). In the extinction phase, the “threat” cue is no longer paired with aversive experiences, with the aim of inhibiting any learned association. In this phase, anxious participants show increased responding to the threat cue, highlighting impoverished threat extinction learning (Duits et al., 2015; Lissek et al., 2005; McGregor et al., 2021).

However, fear conditioning research predominately relies upon descriptive, model-agnostic measures, such as averaged phase responding. These proxies lack explanatory value, mask variance, and inflate results by introducing analytic flexibility (Lonsdorf et al., 2022; Rouder & Haaf, 2019). Computational modelling instead quantifies the hypothesised cognitive-behavioural processes presumed to be generating participant responding (Huys et al., 2016). Reinforcement learning models specifically account for the inherent trial-by-trial threat updating process inherent to fear conditioning (Yamamori & Robinson, 2023). For instance, a lab study individually fitted a reinforcement learning model to physiological responding, revealing a low safety learning rate to be behind the increased responding to the safe cue in an acquisition phase by anxious subjects (Abend et al., 2022). Although typically lab based and with restricted sample sizes, fear conditioning paradigms are increasingly delivered remotely, enabling larger and more diverse samples, at the expense of experimental control and physiological responding (McGregor et al., 2022; Ney et al., 2023). In recent work using a smartphone delivered remote fear conditioning paradigm and self-report responding, this safety learning association was replicated (Kerr et al., 2026). The triangulation of the association across experimental designs and instrumental modalities indicates a stable effect (Treur et al., 2024). This study further demonstrated lower threat extinction learning rates in more anxious participants to be behind increased responding to the threat cue in the extinction phase. Finally, the novel analysis approach exploited intermittent threat cue reinforcement, fitting a threat negative learning rate for non-reinforced threat updates, revealing a third, otherwise hidden association with anxiety severity (Kerr et al., 2026). However, given this was a small, exploratory study, and the first to model remotely collected data, it is unclear whether these parameters and their associations with anxiety would survive replication in a larger, more diverse sample (Gelman & Carlin, 2014; Ney et al., 2018).

Traditional fear conditioning measures have yet to show promise as an anxiety disorder endophenotype, inherent to which would be both individual heritability (larger in magnitude to anxiety symptom heritability), and a clear genetic correlation with anxiety severity (Gottesman & Gould, 2003; Savage et al., 2017). Multiple studies have established moderate univariate heritability estimates of threat responding over several different outcome measure modalities (Box 1) (Hettema et al., 2003; Kastrati et al., 2022; Purves et al., 2021; Savage et al., 2019; Sheerin et al., 2025). Anxiety disorders are themselves moderately heritable, yet to date, only one study has tested for shared genetic influences between anxiety severity and fear conditioning measures, finding no such association (Hettema et al., 2008). However, these studies all used the sub-optimal descriptive, model-agnostic measures typical in fear conditioning, and were further compounded by modest sample sizes and an absence of pre-registered analysis plans, introducing analytic flexibility (Lonsdorf et al., 2017, 2022; Ney et al., 2018). To date, no attempts have been made to establish heritability estimates for computationally modelled cognitive-behavioural mechanisms, a key step in establishing their presumed biological basis.

With the enhanced statistical power facilitated by remote collection, a large independent sample was previously collected in a nationwide twin cohort, presenting a unique opportunity to explore a novel anxiety disorder endophenotype. Through a pre-registered analytic plan and set of hypotheses, this study first attempted to directly replicate the computational model fitting, and subsequent associations seen between anxiety severity, and threat negative, safety, and threat extinction learning rates (Kerr et al., 2026). Second, univariate and bivariate twin modelling was used to quantify the heritability of learning rates, and any genetic or environmental correlations with anxiety severity. Combined, these experiments would establish whether computationally modelled threat learning rates represent a reliable anxiety disorder endophenotype (Karvelis et al., 2023).

**Box 1.**
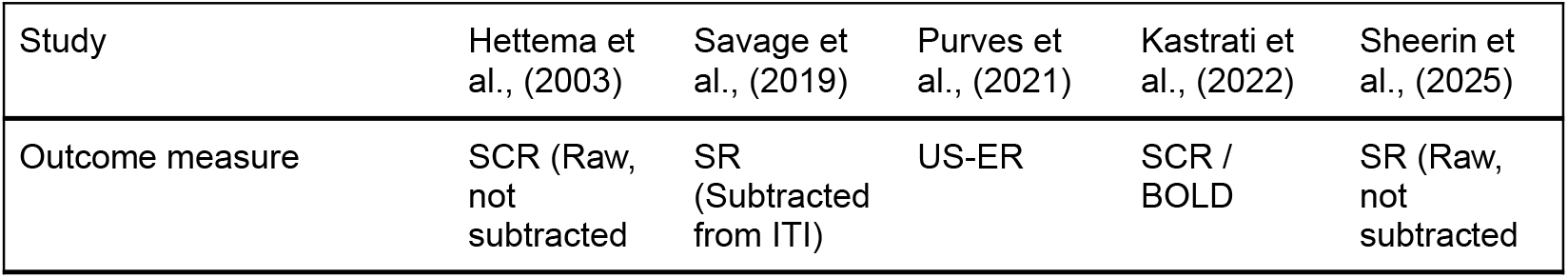
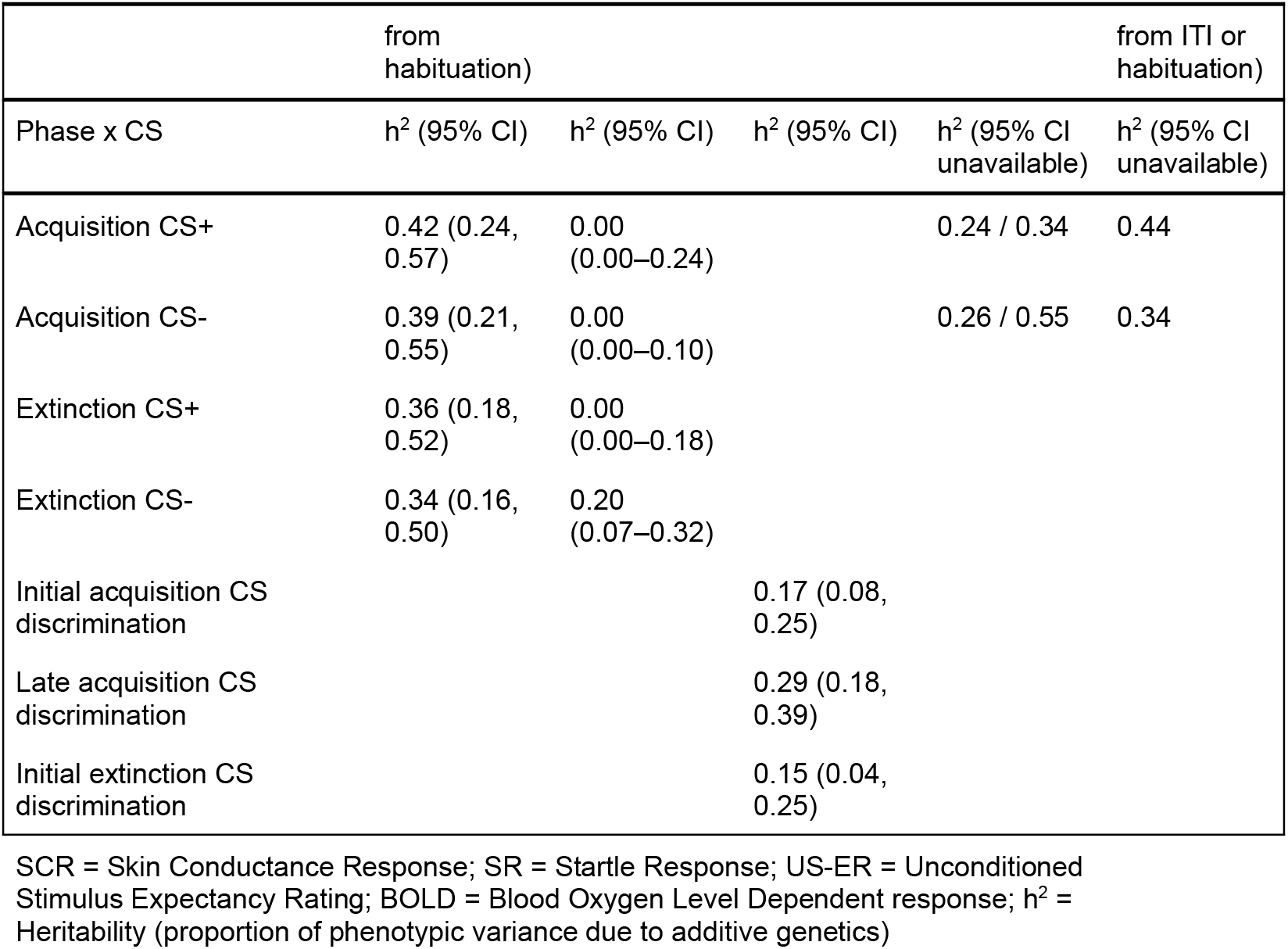

## Results

### Replication

#### Sample & task description

A total of 2575 participants, drawn from a large twin cohort study, completed a remotely delivered differential fear conditioning paradigm, described in previous work and fully in the methods section (Figure 1) (McGregor et al., 2021; Purves et al., 2021). Between phases, participants completed an anxiety severity measure (GAD-7), and a depression severity measure (PHQ-8). Pre-registered “strict” post-task exclusion criteria were applied to ensure full task adherence, leaving 1151 participants from 925 families (Table 2).

**Figure 1.**
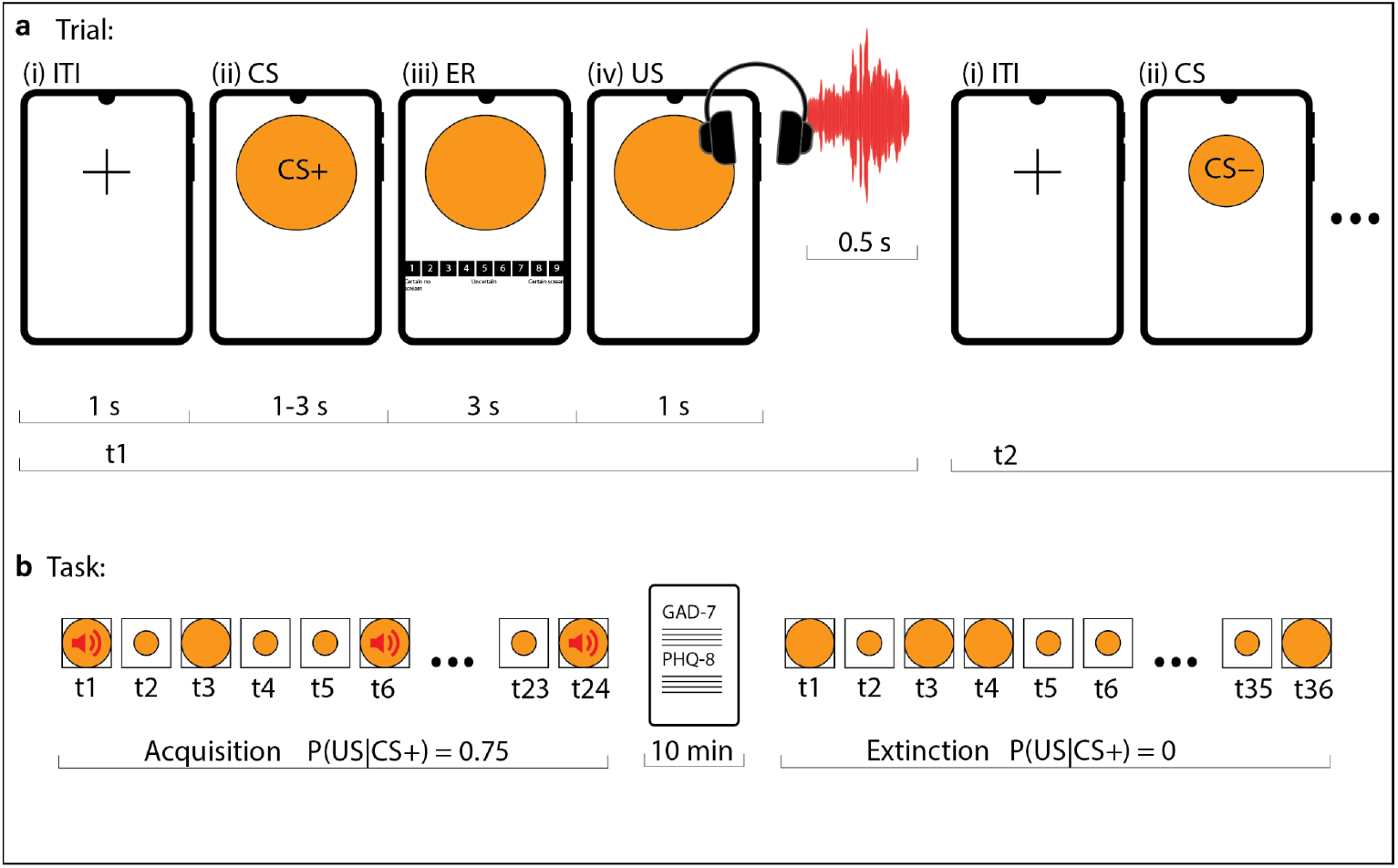
Smartphone delivered sequential trial by trial fear conditioning paradigm. a) Following a fixation cross used as the intertrial interval (ITI) (i), participants were presented with one of two Conditioned Stimuli (CS) (ii). Next, they were presented with an Unconditioned Stimulus (US) Expectancy Rating (ER) scale (iii), upon which they entered their subjective rating of the likelihood of the US occurring. If this trial was a reinforced CS+ trial, US was delivered following the rating, a loud scream noise played through headphones at maximum volume (iv). The next trial began following an ITI. b) The fear conditioning task consisted of an acquisition phase, containing twelve CS+ trials and twelve CS− trials. CS+ was reinforced on 75% of trials. An extinction phase followed, consisting of eighteen CS+ and eighteen CS− trials, neither with reinforcement. Between phases, participants completed the psychiatric measures GAD-7 and PHQ-8.

#### Five learning rate model was best fitting

Computational learning models were applied to the fear conditioning data. Directly replicating previous work, seven Rescorla-Wagner models were tested, featuring differing combinations of learning rates (Table 1) (Kerr et al., 2026). Hierarchical Bayesian modelling was implemented in Stan using Markov Chain Monte Carlo (MCMC) sampling to estimate subject level learning rate parameters. All seven models ran successfully, achieving adequate chain mixing (*r-hat* ≤ 1.01), with no divergences or warnings.

**Table 1.**
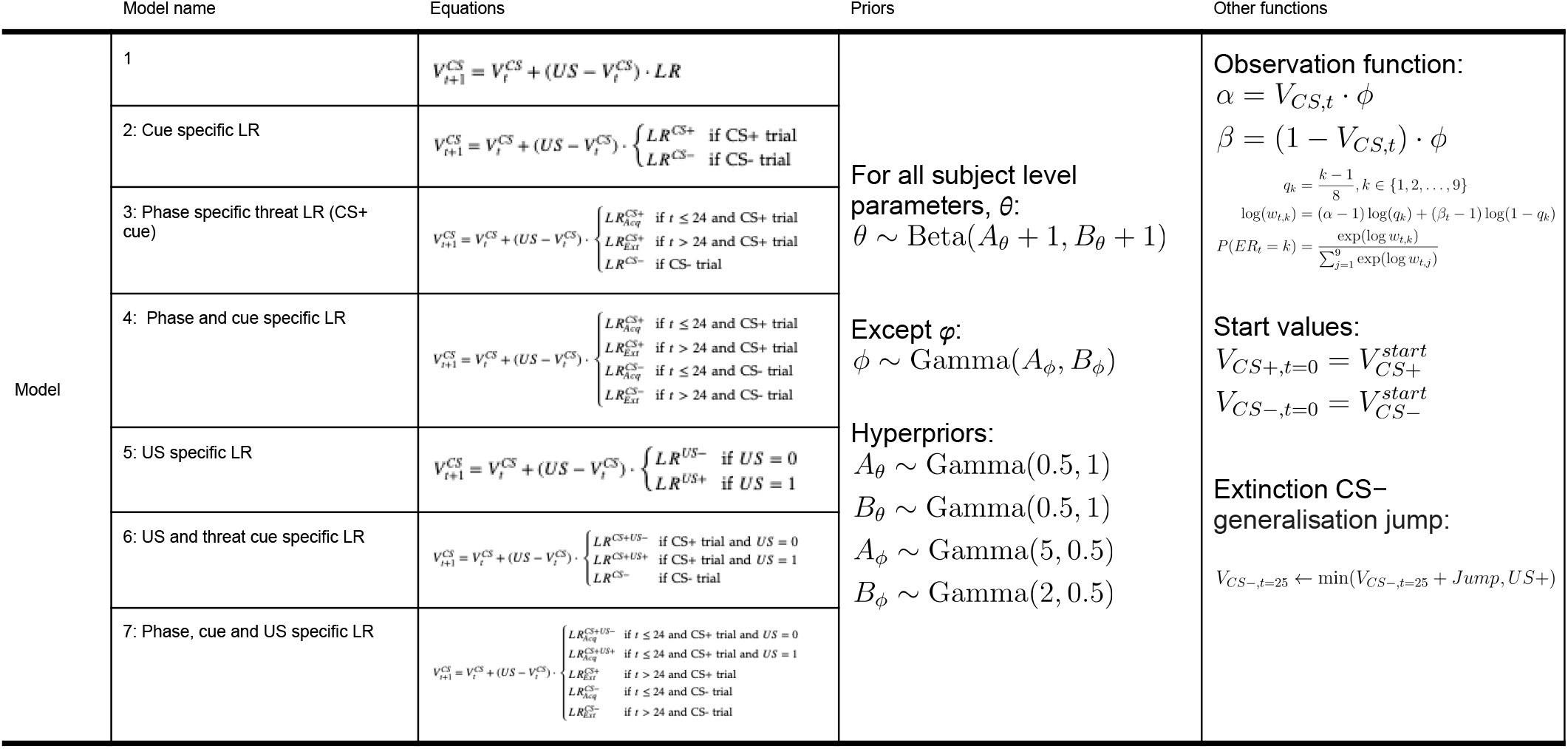
Model specification.

**Table 2.** Participant demographics compared across exclusion criteria groups. Values denote mean or median values with standard deviation (SD) or interquartile range (IQR). Chi-square test used for sex; Wilcoxon rank-sum test used for categorical variables (age, measures). No significant differences in demographics nor anxiety and depression measures were noted between groups. Δn = difference in sample size between moderate and strict exclusion criteria; GAD-7 = Generalised Anxiety Disorder 7 item measure; PHQ-8 = Patient Health Questionnaire 8 item measure.

| Exclusion criteria | n | Female n (%) | Age mean (SD) | GAD-7 median (IQR) | PHQ-8 median (IQR) |
| --- | --- | --- | --- | --- | --- |
| Strict exclusion | 925 | 639 (69.1%) | 23.7 (0.9) | 4 (7) | 4 (6) |
| Moderate exclusion | 1059 | 734 (69.3%) | 23.7 (0.9) | 4 (7) | 4 (6.5) |
| Comparison (Strict vs Moderate) | $\Delta n = 134$ | $\chi^2=0, p=0.951$ | $W=491683.5, p=0.882$ | $W=478990.5, p=0.394$ | $W=483255, p=0.607$ |

In the first of five pre-registered hypothesis sets (*H*_*1*_), using fixed effects Bayesian model selection, we confirmed that a model with five learning rates was best fitting (Figure 2a, Supplementary table 2). This model fitted three learning rates towards the threat cue (CS+); two in the acquisition phase (one for reinforced trials, threat positive learning; another for non-reinforced trials, threat negative learning), and a third for the threat cue in extinction phase (threat extinction learning). Two learning rates were fitted for the safety cue (CS−); one in the acquisition phase (safety learning), the other in the extinction phase (safety confirmation learning). A sensitivity analysis confirmed this model to be best fitting when examining model fit in each phase and conditioned stimulus cue (CS) in isolation (Supplementary table 3).

**Figure 2.**
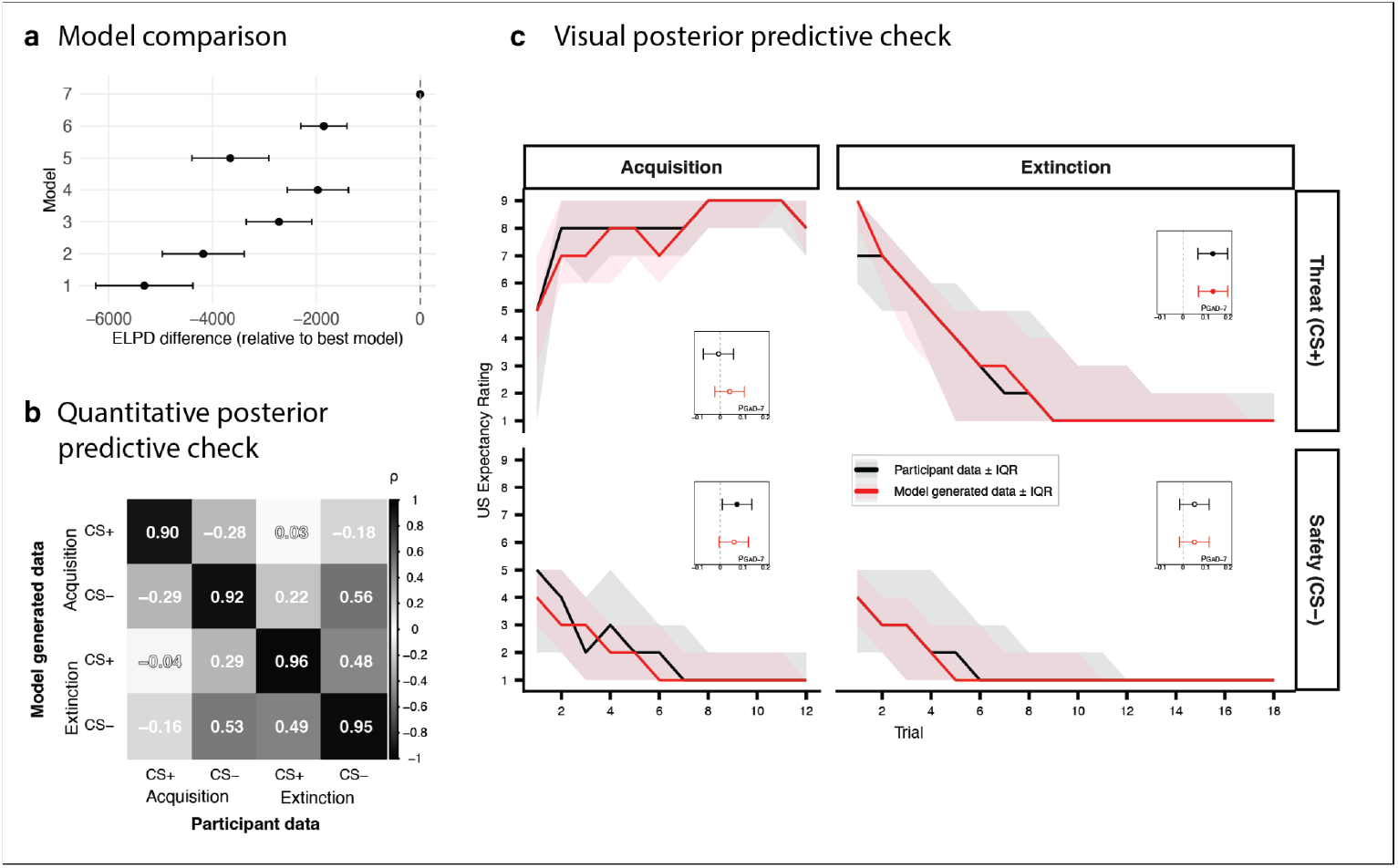
Model comparison and posterior predictive checks. a) The difference in expected log predictive density (ELPD) for each model relative to the winning model, 7, which was significantly better fitting than all others. Error bars signify five times the standard error of the difference, far more conservative than the typical 1.96 critical value for 95% confidence intervals. b) The winning model was used to generate means of whole phase responding, which were compared to participant data equivalents. High Spearman correlations noted across all four phases and CS indicated the model was correctly identifying individual differences in responding. c) Posterior predictive check, with model generated responding overlaid on participant responding. Lines denote trial by trial median responding, and shaded ribbons interquartile ranges (IQR). A further quantitative test of model fit is shown via the inset forest plots. The model generated, and participant, whole phase means from (b) were correlated to GAD-7, our anxiety severity measure of interest. Points indicate Spearman correlation coefficient, with error bars representing 95% bootstrapped confidence intervals. The model successfully replicated an association in extinction phase, however failed to replicate the small association in acquisition phase, albeit the association is on the fringes of significance and the effect sizes are very similar.

In a series of posterior predictive checks, four model-generated descriptive measures (the mean of whole phase responding towards each cue) were compared to their participant generated equivalents (Figure 2b). Spearman correlations between each were marginally higher in extinction phase (*ρCS+* = 0.96 & *ρCS*− = 0.95) than acquisition phase (*ρCS+* = 0.90 & *ρCS*− = 0.92). Next, to assess empirical validity, associations between the model generated descriptive measures and GAD-7 anxiety severity were compared to their participant counterparts (Figure 2c, Supplementary table 4) (Palminteri et al., 2017; Wilson & Collins, 2019). The model reproduced a significant association between GAD-7 and extinction CS+ mean responding (*ρ*_*real*_ = 0.13 [0.07, 0.20] & *ρ*_model_ = 0.13 [0.07, 0.20]), however it did not reproduce a significant association in acquisition CS− (*ρ*_*real*_ = 0.07 [0.01, 0.14] & *ρ*_*model*_ = 0.06, [−0.01, 0.13]).

### Association between anxiety severity and threat extinction learning successfully replicated, but not threat negative nor safety learning

We assessed the replicability and precision of previously established associations between learning rates and anxiety or depression symptom severity through a second pre-registered hypothesis set (*H*_*2*_) (Table 3, Figure 3a(i)). Median point estimates of subject level learning rates were extracted from the model fit and used as dependent variables. As predicted, the threat extinction learning rate demonstrated a small significant negative association with GAD-7 anxiety severity (*ρ* = −0.14 [−0.21, −0.08]). Accordingly, the replication Bayes factor was *BF*_*r*0_ = 1189. 67, indicating the association was over one thousand times more likely under the original study’s finding than under a null model (Figure 3a(ii), Supplementary table 7).

**Figure 3.**
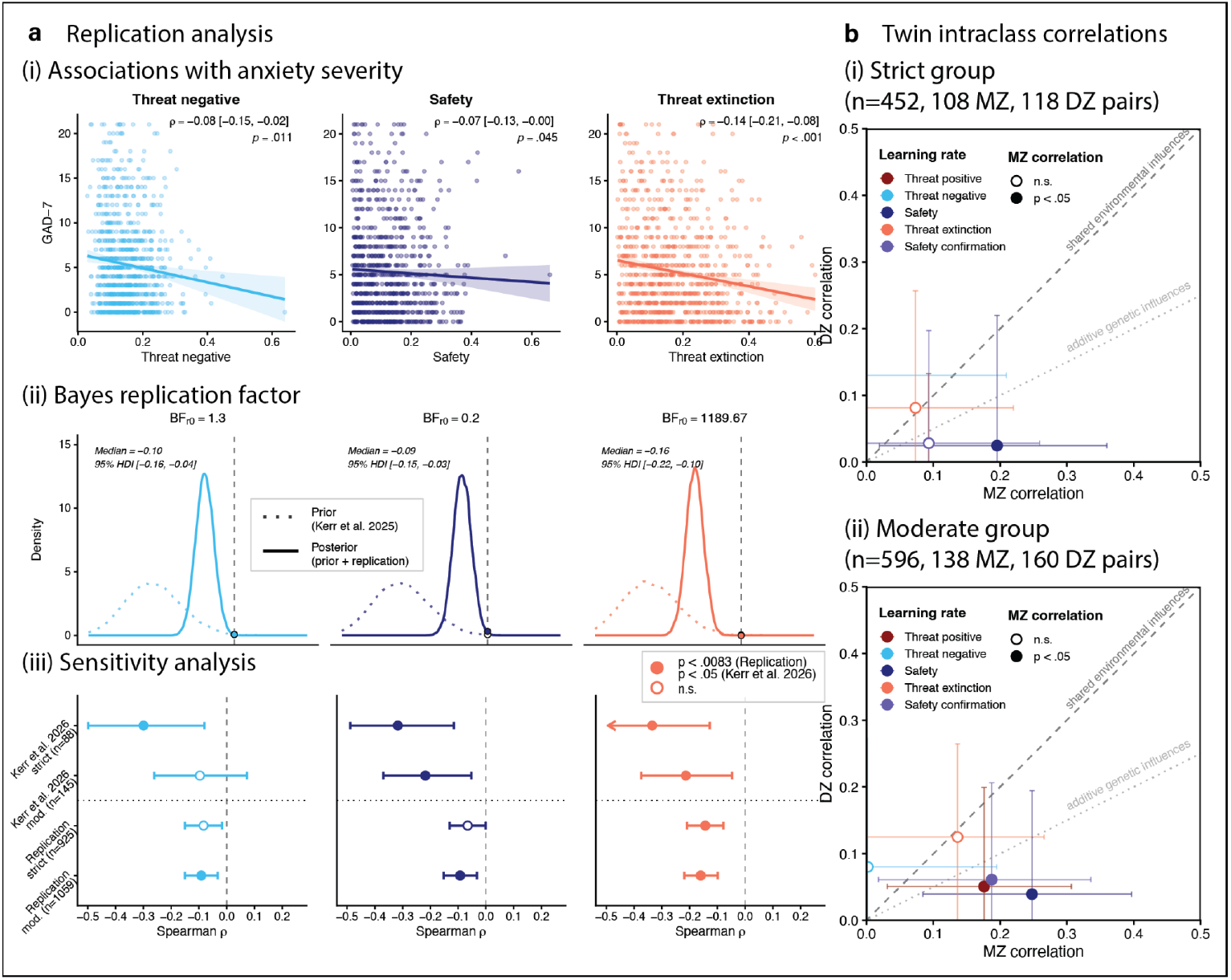
Replication and heritability analyses. a) i) Scatter plots with Spearman correlations of three learning rates previously shown to have associations with GAD-7 anxiety severity, per Kerr et al. (2026). Line of best fit from a linear regression is provided with the shaded ribbons indicating 95% confidence intervals of the regression estimate. ii) Savage-Dickey density ratio plots, visually demonstrating replication Bayes factors (*BF*_*r*0_). The replication Bayes factor is derived from the ratio of the height of the dotted line (the prior, from Kerr et al. (2026)) to the solid line (the posterior) at the point of the null hypothesis (ρ = 0). iii) Forest plot of the sensitivity analysis conducted on exclusion criteria. Points indicate Spearman correlation coefficient, with error bars representing 95% bootstrapped confidence intervals. In Kerr et al. (2026), the strict exclusion group demonstrated larger correlation strengths, however in the replication, the exclusion criteria made minimal difference to correlation strength. b) Scatter plots comparing monozygotic (MZ) to dizygotic (DZ) twin correlations, where greater MZ correlations tend to signify greater genetic influences on a trait. Points represent Intraclass Correlation Coefficient (ICC), with error bars representing 95% confidence intervals. i) In the strict group, only the safety learning rate had a significant MZ correlation. ii) In the moderate group, threat positive and safety confirmation also demonstrated significant MZ correlations. Threat extinction, despite having the strongest association to anxiety severity, was not shown to have genetic influences.

**Table 3.** Spearman correlation testing between each computationally modelled learning rate, and the GAD-7 and PHQ-8 measures of anxiety and depression severity. The analysis was performed in the strict exclusion criteria group (n=925), to test hypothesis set *H2*, before being repeated in the moderate group (n=1059) for a sensitivity analysis, per hypothesis set *H3*. Values are spearman correlation coefficients (ρ) with bootstrapped 95% confidence intervals (CI) in brackets. Bold highlighted values denote a statistically significant correlation, surviving a post-hoc Bonferroni correction (α = 0.05/6) for multiple comparisons, for our six tests per hypothesis set. GAD-7 = Generalised Anxiety Disorder 7 item measure; PHQ-8 = Patient Health Questionnaire 8 item measure.

| Learning Rate | GAD-7 |  | PHQ-8 |  |
| --- | --- | --- | --- | --- |
|  | n = 925 | n = 1059 | n = 925 | n = 1059 |
| Threat positive | -0.06 [-0.13, 0.00] | **-0.11 [-0.17, -0.05]** | -0.03 [-0.09, 0.04] | -0.07 [-0.13, -0.01] |
| Threat negative | -0.08 [-0.15, -0.02] | **-0.09 [-0.15, -0.03]** | -0.06 [-0.13, 0.00] | -0.06 [-0.12, 0.00] |
| Safety | -0.07 [-0.13, -0.00] | **-0.09 [-0.15, -0.03]** | -0.08 [-0.15, -0.02] | **-0.08 [-0.14, -0.02]** |
| Threat extinction | **-0.14 [-0.21, -0.08]** | **-0.16 [-0.22, -0.10]** | **-0.14 [-0.21, -0.07]** | **-0.13 [-0.19, -0.07]** |
| Safety confirmation | **-0.12 [-0.18, -0.05]** | **-0.10 [-0.17, -0.04]** | **-0.10 [-0.17, -0.04]** | **-0.10 [-0.16, -0.04]** |

In contrast, neither the safety learning rate, nor the threat negative learning rate were significantly associated with GAD-7 anxiety severity when corrected for multiple comparisons, with replication Bayes factors of *BF*_*r*0_ = 0. 20 and *BF*_*r*0_ = 1. 30 respectively.

A similar pattern emerged for PHQ-8 depression severity, with the threat extinction learning rate showing a significant negative association (*ρ* = −0.14 [−0.21, −0.07]), and, as predicted, no significant associations for the safety learning rate, and threat negative learning rate. Finally, the residual scores for GAD-7 and PHQ-8 failed to display significant associations for any learning rate, with the associations fully accounted for by shared variance between GAD-7 and PHQ-8 (Supplementary table 5).

Whilst not within our formal hypothesis testing framework, a significant negative association was noted between the safety confirmation learning rate and both GAD-7 and PHQ-8 (*ρ* = −0.12 [−0.18, −0.05] & *ρ* = −0.10 [−0.17, −0.04]).

#### Poorer task adherence increases associative effect sizes

In a sensitivity analysis, a third pre-registered hypothesis set (*H*_*3*_) tested the impact of task adherence on association effect size, in which poorer adherence had previously attenuated effect size. From the total of 2575 participants, 1357 participants survived the pre-registered “moderate” exclusion criteria, containing 1059 unique families. In this group, participants who reduced their headphone volume to 80% of the maximum were included, and contingency awareness was not mandated (Supplementary table 1). The demographics of this group, and distributions of our measures, did not differ significantly to the “strict” group (Table 2). Posterior predictive checks of model fitting were similar; the significant association between GAD-7 and descriptive extinction CS+ mean responding was again reproduced by the model (*ρ*_*real*_ = 0.13 [0.07, 0.19] & *ρ*_model_ = 0.13 [0.07, 0.19]), however, now, the significant association to acquisition CS− mean responding was also reproduced (*ρ*_*real*_ = 0.11 [0.05, 0.17] & *ρ*_model_ = 0.10 [0.03, 0.16]) (Supplementary Table 4).

We predicted that within this larger, less rigorous sample, the threat extinction learning rate would show a reduced association magnitude with GAD-7 anxiety severity compared to the strict sample, however, the significant negative association was marginally stronger (*ρ* = −0.16 [−0.22, −0.10]), with a much larger replication Bayes factor (*BF*_*r*0_ = 283655. 07) (Figure 3a(iii), Supplementary tables 6 & 8). Similarly against prediction, this sample now displayed a significant negative association between safety learning and GAD-7 anxiety (*ρ* = −0.09 [−0.15, −0.03]). Also against prediction, the threat negative learning demonstrated a significant negative association with GAD-7 anxiety (*ρ* = −0.09 [−0.15, −0.03]), despite its absence in the original study.

The threat extinction learning rate maintained a significant negative association with PHQ-8 (*ρ* = −0.13 [−0.19, −0.07]), and was as predicted marginally reduced in magnitude to that from the strict exclusions sample. Against prediction, the safety learning rate demonstrated a significant negative association with PHQ-8 (*ρ* = −0.08 [−0.14, −0.02]), whereas, with prediction, the threat negative learning rate did not. Finally, decomposing the association between threat extinction learning and GAD-7 demonstrated significant negative associations with both shared and unique variance to GAD-7 (*ρ*_*shared*_ *=* −0.15 [−0.21, −0.09 & *ρ*_*unique*_ = −0.10 [−0.16, −0.04]) (Supplementary table 6).

Also outside of our formal hypothesis testing framework, mirroring the strict sample, a significant negative association was noted between the safety confirmation learning rate and both GAD-7 and PHQ-8 (*ρ* = −0.10 [−0.17, −0.04] & *ρ* = −0.10 [−0.16, −0.04]). Further, the threat positive learning rate demonstrated a significant negative association to GAD-7 alone (*ρ* = −0.11 [−0.17, −0.05]).

### Heritability

#### Only the safety learning rate is heritable

By comparing the similarity of monozygotic (MZ) twins, who share all segregating alleles, to dizygotic (DZ) twins, who share approximately half, one can estimate the relative influences of genetics and environment on phenotypes, including learning rates. Heritability is roughly estimated as twice the difference between the MZ and DZ correlations (which estimate their degree of similarity) for any trait. We predicted moderate heritability estimates (0.3 < *h*^*2*^ < 0.5) for all five learning rates (*H*_*4*_) in the “strict” exclusions group, containing 108 MZ, and 118 DZ pairs.

The best fitting computational model was refitted to the sample of complete twin pairs meeting the strict exclusion criteria, with median point estimates of subject level learning parameters extracted from the fit. Intraclass correlations, which control for any birth order effects, were calculated for each learning rate parameter. However, in this strict exclusions sample, only the safety learning rate demonstrated a significant MZ correlation (*r*_*MZ*_ = 0.20 [0.02, 0.36]), with all others non-significant, and often lower than DZ correlations, indicative of low statistical power (Figure 3b, Table 4).

**Table 4.**
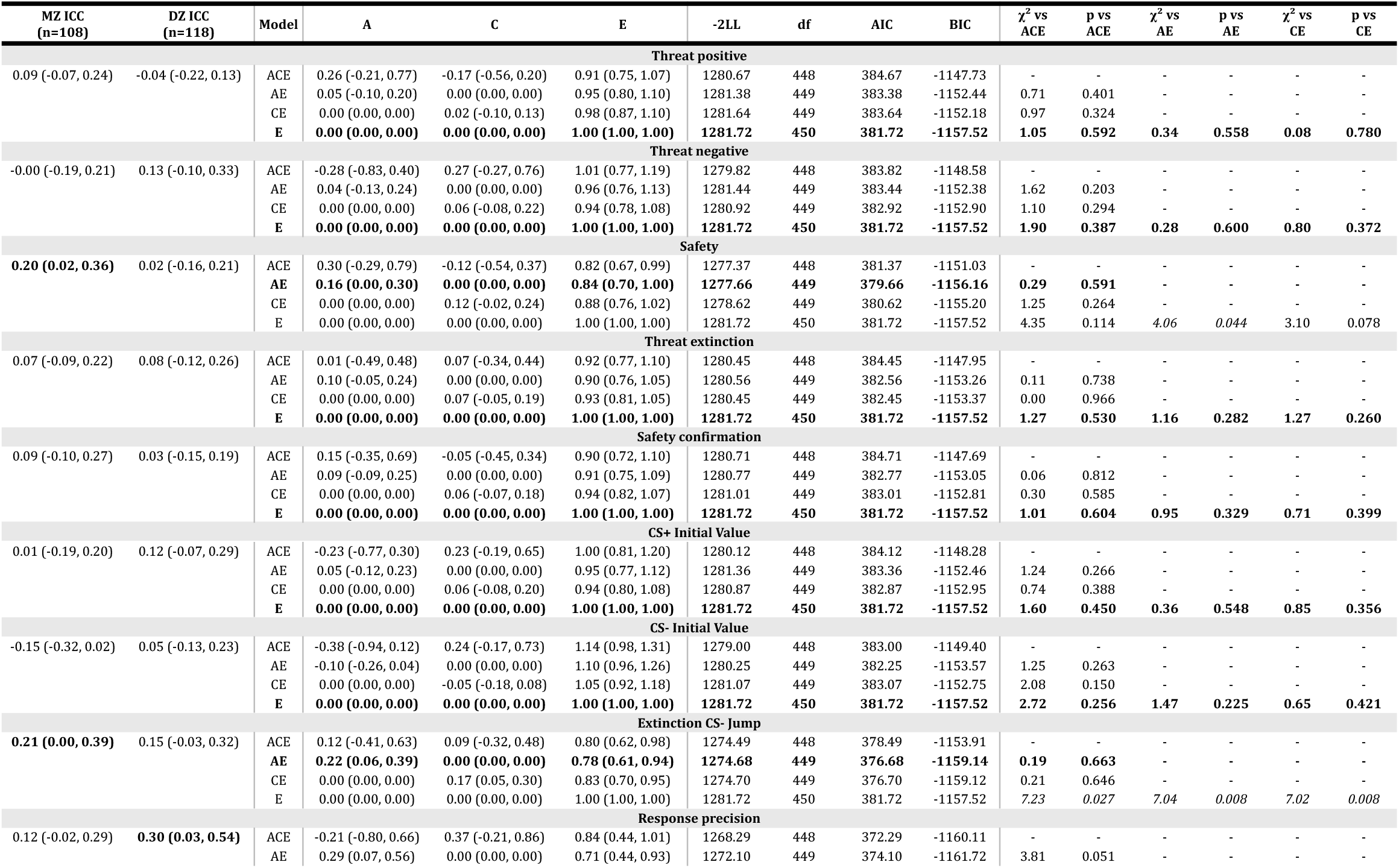
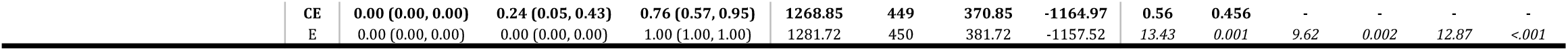
Full univariate heritability testing of computationally modelled learning rates, incorporating twin correlations (intraclass correlation coefficient, ICC), and full ACE model fitting. For ICC testing, 95% confidence intervals (CI) were calculated via bootstrapping (10,000 iterations); bold highlighted values indicate a significant ICC value with 95% CI excluding 0. Differences in twin correlations (with MZ > DZ) indicate increased genetic influences, whereas similar correlations (MZ = DZ) indicate greater shared environmental influences (developmental or rearing effects). For ACE model fitting, ACE twin models were fitted and compared to models with factors dropped, with the best fitting model not significantly worse fitting than ACE (italic χ^2^ value indicates significantly worse fit), and with the lower AIC in the event of a tie between AE and CE models. Bold highlighted rows indicate the best fitting model per hierarchical likelihood testing; 95% confidence intervals for parameter estimates were calculated via bootstrapping (1000 iterations). A = additive genetic; C = shared environment; E = unique environment + error. The safety learning rate was the only learning rate to display heritability.

To formally quantify heritability, univariate ACE modelling was applied to all learning parameters, to decompose phenotypic variance into three components; shared (*C*) and non-shared (*E*) environmental, and additive genetic factors (*A*), synonymous with heritability (*h*^*2*^). The best fitting model for the safety learning rate was the AE model (dropping the *C* factor), with a significant heritability estimate (*h*^*2*^_*safety*_ = 0.16[0.00.0.30]) (Table 4). Given the non-significant MZ and DZ twin correlations, the best fitting model for the remaining learning rates was the E model, dropping both *A* and *C* components, precluding heritability estimates.

#### Exploratory analysis of univariate heritability by increasing power

In our pre-registration, we committed to exploratory analyses should twin models fail. To address low power, we conducted a post-hoc exploratory analysis, using the moderate exclusion criteria sample (consisting of 138 MZ and 160 DZ twin pairs). The two groups did not significantly differ on demographics nor measures tested (Supplementary table 9).

In the moderate exclusions group, significant MZ intraclass correlations were noted within the threat positive learning (*r*_*MZ*_ = 0.18 [0.03, 0.31]), safety learning (*r*_*MZ*_ = 0.25 [0.08, 0.40]), and safety confirmation learning rates (*r*_*MZ*_ = 0.19 [0.02, 0.33]) (Figure 3b, Supplementary table 10). Accordingly, ACE modelling revealed the AE model to be best fitting in each case, with heritability estimates of 16%, 21%, and 17% respectively (*h*^*2*^_*threat positive*_ = 0.16 [0.00,0.30], *h*^*2*^_*safety*_ = 0.21 [0.07, 0.34], *h*^*2*^_*safety confirmation*_ = 0.17 [0.03, 0.32]). Despite non-significant MZ and DZ correlations, the threat extinction learning rate was well fitted by the CE model, dropping the *A* component. This revealed a shared environmental influence of 13% (*C* = 0.13 [0.02, 0.22]).

#### Bivariate heritability

Finally, in accordance with our fifth pre-registered hypothesis, we investigated whether genetic and environmental influences were shared between learning rates and anxiety severity. Cross-twin cross-trait correlations (the correlations between twin 1’s learning rate and twin 2’s anxiety severity, and vice versa) were calculated, before formal quantification using bivariate ACE modelling.

In the strict sample, only threat extinction learning rate had a significant phenotypic correlation to decompose (*r*_*ph*_ = −0.14 [−0.24, −0.05]), and the cross-twin cross-trait correlations between threat extinction learning rate and GAD-7 anxiety severity were small and non-significant suggesting a lack of genetic or shared environmental influences on the association (Supplementary table 11). Similarly, whilst the safety learning rate was heritable, the phenotypic correlation with anxiety was negligible, also resulting in non-significant cross-trait correlations. For these reasons, bivariate modelling was underpowered (Supplementary table 13), resulting in non-significant cross-trait genetic and environmental correlations (*rA* & *rE*) for the safety learning rate (Supplementary table 14). The threat extinction learning rate did however display a significant negative cross-trait environmental correlation (*rE =* −0.17 [−0.32, −0.02]), alongside a non-significant genetic correlation.

In the exploratory moderate sample, the safety learning rate, despite a non-significant phenotypic correlation, displayed a significant cross-twin cross-trait correlation (*r*_*MZ-CTCT*_ = −0.13 [−0.24, −0.03]), whereas the threat extinction learning rate equivalent remained non-significant (Supplementary table 12). Bivariate ACE modelling, where the AE model was best fitting in both cases, similarly resulted in non-significant cross-trait genetic and environmental correlations for safety learning rate, whilst threat extinction learning again displayed a significant negative cross-trait environmental correlation (*rE =* −0.15 [−0.27, −0.01]) (Supplementary table 14).

## Discussion

In this article, we sought to establish the reliability of computationally modelled learning rates as a phenotype in human fear conditioning, the reliability of their associations with anxiety severity, and finally examine their heritability, and potential as an endophenotype.

Firstly, we successfully fitted a five learning rate model, replicating previous work (Kerr et al., 2026). Although the effect size was halved, likely affected by “winner’s curse” inflation in the original study, we further replicated a negative association between threat extinction learning rate and GAD-7 anxiety severity, (*ρ*_original_ = −0.32, *ρ*_replication_ = −0.14) (Button et al., 2013; Ioannidis, 2005). This confirms that those with higher anxiety severity are slower to learn that a once threatening stimuli is now safe, supported by a replication Bayes factor (*BF*_*r*0_ = 1189. 67) indicating extreme evidence of this effect (Stefan et al., 2019). The computational approach provides a reliable mechanistic account of previously model-agnostic, descriptive measures, amplifying individual differences whilst reducing researcher analytic flexibility (Lonsdorf et al., 2022; Rouder & Haaf, 2019). The repeated identification of this association across two independent, heterogenous, remotely collected samples, surviving stringent family-wise error correction, suggests the rate of threat updating is a reliable, robust phenotype of anxiety.

Notably, against our predictions, we failed to replicate two associations from the earlier paper, including safety learning (Acquisition CS-). Impaired safety learning is a robust effect in the literature of model-agnostic fear conditioning measures, albeit likely inflated through publication bias and case-control design (where we are examining dimensional anxiety) (Kausche et al., 2025). However, our sensitivity analysis, with a larger sample owing to more permissive post-task adherence exclusion criteria, did replicate the association (Table 3). The association is present when examined with a descriptive measure, and in another study using descriptive measures and a case-control design within the same cohort (McGregor et al., 2021)). This indicates the task itself is capable of producing individual differences, and this type II error is likely a failure of the computational model (Hedge et al., 2018).

Our quantitative posterior predictive check, the correlations between a model generated and real descriptive measure, were slightly lower in the acquisition phase than extinction, indicating non-modelled variance in the descriptive measure that the model was unable to capture. The Rescorla-Wagner model cannot accede to the sudden counter-factual updating seen in this phase, and, regrettably, variants tested with this feature failed to adequately recover parameters, owing to parameter identifiability issues in minimally featured data (Kerr et al., 2026). Therefore, the safety learning rate is not yet a reliable phenotype of anxiety.

Secondly, examining the genetic basis of each phenotype, we successfully established a modest heritability estimate for the safety learning rate (*h*^*2*^_*safety*_ = 0.16), however we were unable to establish the same for any other learning rate (Table 4). A post-hoc power calculation (albeit one not specifically designed for a repeated measures approach like fear conditioning) indicated a power of 0.20, assuming a true heritability of 0.20. This power deficiency increased the likelihood of non-significant twin correlations and negligible differences between monozygotic and dizygotic twins, likely driving type II error (Button et al., 2013; Krzywinski & Altman, 2013). Our pre-registration detailed our plans to undertake exploratory research in these circumstances, in effect running a further sensitivity analysis against our fourth and fifth hypotheses.

We repeated the heritability analysis in the “moderate” exclusions group, permitting the volume of the aversive stimulus to drop to a minimum of 80%, increasing statistical power to 0.23. This established further heritability estimates for threat positive and safety confirmation learning rates (*h*^*2*^_*threat positive*_ = 0.16, *h*^*2*^_*safety confirmation*_ = 0.17), whilst increasing the estimate for safety learning (*h*^*2*^_*safety*_ = 0.21). Interestingly, the threat extinction learning rate demonstrated significant shared environmental estimates in the moderate group (*C* = 0.13), indicating shared developmental and rearing effects drive this trait most strongly associated with anxiety severity. The threat negative learning rate was always best explained by the saturated (non-shared environment) model, indicating a noisy measure. In summary, we have established empirical evidence of the genetic origins of computationally modelled learning rates, a key step in establishing their biological basis.

Finally, key to our endophenotype proposal was the existence of both a phenotypic correlation (to anxiety severity) and a significant MZ cross-twin cross-trait correlation, suggestive of genetic overlap. Regrettably, whilst we established the former in the threat extinction learning rate, we could not establish the latter in any learning rate. In the moderate group we did establish a modest MZ cross-twin cross-trait correlation for the safety learning rate, raising the possibility of a genetic relationship between safety learning and anxiety severity. Unfortunately, we were unable to confirm this through bivariate modelling.

Whilst low power significantly increases the chance of type II error, given the only previous bivariate study of this kind also failed to produce evidence of pleiotropy, an absence of effect may be more likely than the failure to detect one (Hettema et al., 2008). Given the five previous univariate heritability studies of fear conditioning fail to report an anxiety measure, this ambiguity may be compounded by the file drawer problem (Hettema et al., 2003; Kastrati et al., 2022; Purves et al., 2021; Rosenthal, 1979; Savage et al., 2019; Sheerin et al., 2025). As it stands, the evidence from our study suggests the genetic basis of the safety learning rate is unrelated to the genetic basis of anxiety severity. It is therefore unsuitable as an endophenotype, and as yet unknown developmental effects must contribute to the observed phenotypic association.

The trade off between task adherence and sample size is the key limitation to this work, where despite the large sample size facilitated by the remote delivery of the paradigm, the number of complete twin pairs surviving even the broadest post-task adherence exclusion criteria resulted in a power of only 0.23. We promised additional remuneration for complete twin-pair task completion, which may have increased rates of recruitment of unmotivated second twins via co-twin peer-pressure (66% of the full sample were twin pairs, which dropped to 43% and 39% in moderate and strict exclusions groups). Further, the lack of experimental control, and reliance on self-declarations of compliance, introduces measurement error, only partially offset by the constraining effects of hierarchical Bayesian modelling. The TEDS cohort, whilst similarly sociodemographically and geographically distributed to the UK population (Lockhart et al., 2023), may differ in some aspects to the normative population, for example with reduced birth weights. However, modern studies suggest that any differences have negligible impacts on study outcomes (Christensen & McGue, 2020). Twin studies rely on assumptions which are inevitably not met, including identical shared environmental components (*C*), which are typically more similar in monozygotic twins, inflating heritability estimates (Richardson & Norgate, 2005). Further heritability estimate inflation may arise from gene-environment interactions, which are not directly controllable in our simple univariate ACE models.

In conclusion, we have demonstrated that computationally modelled threat extinction learning rate is robustly associated with anxiety severity, though there was limited evidence for its heritability. Safety learning rate is heritable, providing support for the biological basis of this cognitive-behavioural mechanism. However, we failed to produce any evidence to suggest that either learning rate shares a genetic component with anxiety severity, suggesting they are unsuitable anxiety endophenotypes. Whilst these learning rates remain a key mechanistic feature of anxiety, even if they can be measured with greater reliability in larger samples, they are unlikely to be sufficiently heritable to make them a useful endophenotype through which to explore the genetics of anxiety disorders.

## Online methods

### Ethical approval

Ethical approval was granted by the King’s College London Psychiatry, Midwifery and Nursing Research Ethics Subcommittees (application PNM/09/10-104)

### Pre-registration

The study was pre-registered on OSF (https://osf.io/7b2d8/overview), containing the five sets of hypotheses tested in this paper. The first three pertained to the replication analysis (*H*_*1*_, *H*_*2*_, *H*_*3*_). The latter two, *H*_*4*_ and *H*_*5*_, pertained to the later heritability analysis.

### Replication

#### Sample

The sample was drawn from the Twin Early Development Study (TEDS), a longitudinal twin study of re-contactable monozygotic (MZ), dizygotic (DZ) twin pairs. Of the 5,914 participants contacted, 2,987 responded and consented, with 2925 passing an initial screening phase (excluding those with a past medical history of heart, neurological, or auditory pathology, or currently pregnant). Of these, 2575 completed the task up to the end of the extinction phase. Twins were paid £10 for completion, with the added incentive of inclusion on prize draws if both twins completed the task.

#### Task

A fear conditioning paradigm was delivered remotely via the FLARe smartphone app (McGregor et al., 2022), as previously published in (McGregor et al., 2021; Purves et al., 2021). In brief, the paradigm delivered two phases; acquisition and extinction. Each phase consisted of repeated presentations of two conditional stimuli (CS+ and CS-; different sized coloured circles). The CS+ was paired with an aversive unconditional stimulus (US; a loud scream noise) on 75% of acquisition trials only. The extinction phase had no US pairing. Participants rated their expectancy of the US at each CS presentation (US-ER), on a 9 point Likert scale (Figure 1).

#### Task exclusions

Compliance with the task was assessed via app generated data and self-report questions which assessed if participants followed instructions, removed headphones, altered the volume, restarted the app, whilst also testing participants’ perception of US unpleasantness, and contingency awareness.

Pre-registered task exclusion criteria were applied in two phases, a strict criteria for the replication study, a moderate criteria for the replication study sensitivity analysis, and, as it would transpire, for exploratory heritability analyses (Supplementary Table 1). The demographics of these two samples were compared for significant differences in age, sex, or psychiatric measures used in the paper (Table 2). Given the assumption of statistical independence between participants in correlation testing, no complete twin pairs were used in the replication analyses. Following exclusion criteria application, all singletons (unmatched twins) and one randomly selected twin from each complete pair were retained for subsequent model fitting and analysis.

#### Measures

In the break between fear conditioning phases, participants completed the Generalized Anxiety Disorder Assessment (GAD-7) and Patient Health Questionnaire-8 (PHQ-8) via the app interface.

#### Computational modelling

The modelling process was a direct replication of (Kerr et al., 2026). As per the pre-registration, seven Rescorla-Wagner reinforcement learning models were fitted to trial-by-trial US-ER data. Models varied in the number of, and structure of, learning rate parameters. Full model specifications are listed in Table 1.

For each trial, the models updated a latent associative value assigned to trials of each CS (*V*_*CS+*_, *V*_*CS*−_) through prediction error learning, weighted by an individual learning rate. To map this continuous value to the ordinal 9 item US-Expectancy Rating scale, a discretised beta distribution observation function was used. At the beginning of each trial, the continuous estimated associative value, *V*, was transformed into a beta distribution, with *V* acting as the mean, and the width set by a response precision parameter, phi (*φ*).

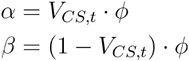

This distribution was evaluated at nine proportional anchor points (*q*_*k*_), to produce a weight for each response (*w*_*k*_), which was passed through a categorical softmax function to calculate the likelihood of the US-Expectancy Rating of that trial (*ER*_*t*_). In the event of a missed trial, the likelihood evaluation and value update were skipped for that trial.

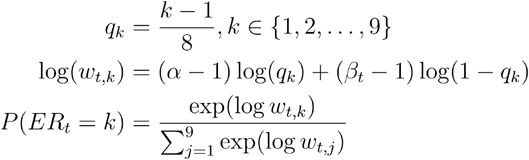

The associative value, *V*, was updated via a Rescrola-Wagner learning model, with prediction error updating weighted by a subject specific fitted learning rate. The prediction error was driven by the presence or absence of *US* (*US+* = 1, *US*− = 0). In the base model (model 1), a single learning rate is used for all trials.

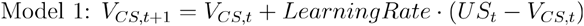

In subsequent models, learning rates are separated into one per CS (model 2), one per phase of CS+ trials, and a third for all CS− trials (model 3), and separate learning rates for each phase and CS (model 4).

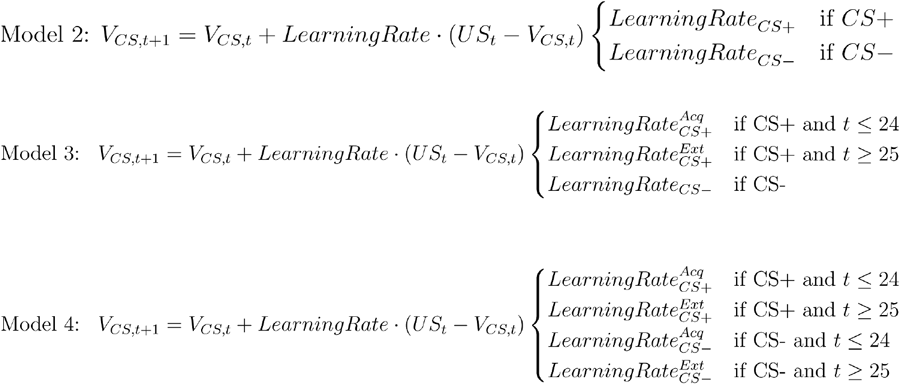

A further three models incorporated differential learning to positive and negative outcomes. This learning was only possible in CS+ Acquisition trials, where *US* differed. Model 5 incorporated two learning rates, one updating for positive trials (*US+*), and the other for negative trials (*US*−). Model 6 fitted a third learning rate for all CS− trials (effectively testing if the negative updates in Acquisition CS+ are subject to the same learning process as the Extinction phase). Finally, model 7 fitted separate learning rates to all phases and CS, in addition to the two learning rates for Acquisition CS+ trials.

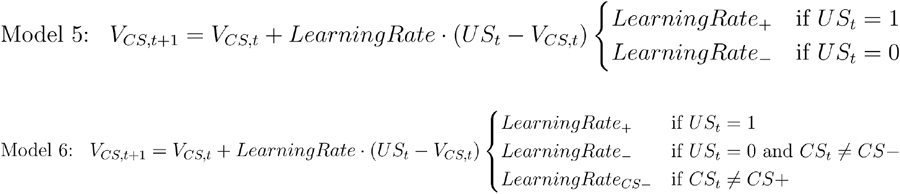

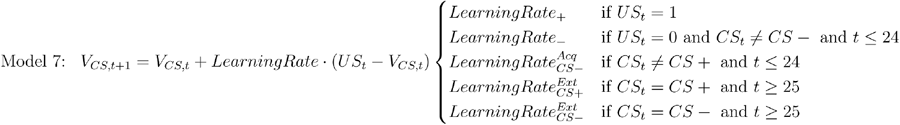

Both starting values (*V*_*CS+,t=0*_, *V*_*CS*−,*t=0*_) were free parameters fitted by the model. A final CS− generalisation “jump” parameter was also fitted, which increased *V*_*CS*−_ between the Acquisition and Extinction phase.

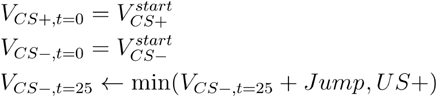

All model parameters were estimated in a hierarchical Bayesian framework, with group level hyperparameters constraining subject level parameters. If *θ* represents any bounded subject level parameter (learning rates, start values, CS− generalisation jump), *A*_*θ*_ and *B*_*θ*_ were group level hyperparameters within a beta distribution, both iterated by 1 to ensure a unimodal distribution.

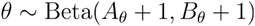

The positive unbounded phi parameter, *φ*, was instead drawn from a gamma distribution, similarly with two hyperparameters *A*_*φ*_ and *B*_*φ*_.

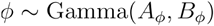

Broad, uninformative priors were used, with the exception of the phi parameter, *φ*, which had a stronger priors to reduce the likelihood of extreme parameter estimates.

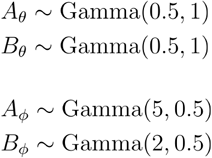

All seven models were fit with 2000 warm-up and 4000 sampling Markov Chain Monte Carlo (MCMC) iterations, using four parallel chains. Models were implemented in Stan (CmdStan version 2.37.0) and run within R scripts using the package cmdstanr (version 0.9.0) on instances of R (version 4.5.1) running on the Create High Performance Cluster at KCL (King’s College London e-Research team, 2022). Given the sample size, within-chain parallelisation was implemented using the reduce_sum function within Stan, with the grainsize argument applied to subjects, where eight cores were allocated per chain. Following fitting, models were checked for divergences using the cmdstan_diagnose function. Chain convergence was assessed via Gelman-Rubin r-hat value, 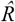, with 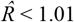 considered acceptable (Baribault & Collins, 2023). Qualitative posterior predictive checks were performed at group level to visually confirm adequate model fit.

To select the best fitting model, fixed effects Bayesian model selection was performed. Using the loo_compare function from the loo package (version 2.9.0), the expected log predictive density (ELPD) of each model was calculated. The best model was determined as having the highest ELPD, with five times the standard error of the difference not exceeding the difference to the next best model. Further, a pseudo-*R*^*2*^ value was calculated, to compare each model’s likelihood, *L*_ℳ_ to chance responding, *L*_𝒞_.

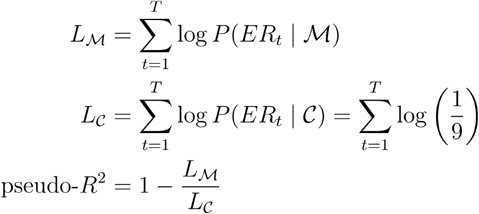

Our first hypothesis, *H*_*1*_, was: Model 7 will be the best fitting model, determined by the lowest ELPD, containing five learning rate parameters (threat positive learning rate, threat negative learning rate, threat extinction learning rate, safety learning rate, and safety confirmation learning rate).

This model selection process was repeated for each phase and CS separately, to examine for model bias and underfitting. Median point estimates of the winning model’s learning rate(s) were extracted from the model fit, and used as dependent variables in the subsequent analyses.

#### Associations

To examine associations between learning rates (from the winning model) and anxiety and depression severity, through the measures GAD-7 and PHQ-8, non-parametric Spearman’s rho correlations were calculated (given GAD-7 and PHQ-8 are non-normally distributed). For each learning rate, 95% confidence intervals were estimated using bootstrapped sampling with 10,000 iterations. Per the pre-registration, a bonferroni correction was applied for the six hypothesis tests performed 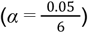, listed below.

*H2*_*a*_: Safety learning rate will display a negative association with anxiety severity.

*H2*_*b*_: Threat extinction learning rate will display a negative association with anxiety severity.

*H2*_*c*_: Threat negative learning rate will display a negative association with anxiety severity.

*H2*_*d*_: Safety learning rate will display no association with depression severity.

*H2*_*e*_: Threat extinction learning rate will display a negative association with depression severity.

*H2*_*f*_: Threat negative learning rate will display no association with depression severity.

Given the high correlation between GAD-7 and PHQ-8, we computed a rank based residualisation analysis. Shared variance was calculated as the mean of z-scored ranks. Unique variance was calculated as the residual from a linear model predicting the rank of one measure from the rank of the other. These components were correlated to each learning rate using Spearman’s rho and bootstrapped 95% confidence intervals with 10,000 iterations.

To assess the impact of task adherence dictated by our exclusion criteria, a sensitivity analysis repeated these steps using the moderate exclusion criteria sample. The pre-registered hypotheses are listed below.

*H3*_*a*_: Safety learning rate will display a weaker negative association with anxiety severity than in the strict exclusion sample.

*H3*_*b*_: Threat extinction learning rate will display a weaker negative association with anxiety severity than in the strict exclusion sample.

*H3*_*c*_: Threat negative learning rate will display no association with anxiety severity.

*H3*_*d*_: Safety learning rate will display no association with depression severity.

*H3*_*e*_: Threat extinction learning rate will display a weaker association with depression severity than in the strict exclusion sample.

*H3*_*f*_: Threat negative learning rate will display no association with depression severity.

#### Bayes replication factor analysis

To examine the success or failure of a replication of effects from Kerr et al. (2026), a Bayesian replication factor was calculated for each learning rate association tested, which computes the probability of the replication effect given the original study effect (*BF*_10_(*d*_*rep*_ |*d*_*orig*_ = *BF*_*r*0_).

Using an evidence updating framework, a replication hypothesis (*H*) informed by the posterior distribution from Kerr et al. (2026), was tested against a null hypothesis (*H*_0_), containing no prior information.

Given the non-parametric Spearman associations used, variables were ranked and standardised within each sample independently, to prevent between study differences in scale or location impacting the pooling (Ly et al. 2019). Correlation parameter estimates (median ρ) were derived from MCMC sampling via the BayesFactor package (version 0.9.12-4.7), using 20,000 sampling iterations. 95% High Density Intervals were calculated using the coda package (version 0.19-4.1). Bayes factors were calculated using the BayesFactor package, using the default Jeffreys-Zellner-Siow (JZS) prior of 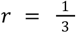, in keeping with our correlation effect sizes from Kerr et al. (2026). Bayes factors were calculated for the associations from Kerr et al. (2026) (*BF*_10_(*d*_*orig*_) = *BF*_*orig*_), before the data were pooled, and a pooled Bayes factor calculated (*BF*_10_(*d*_*orig*_, *d*_*rep*_) = *BF*_*pooled*_). The replication Bayes factor was derived by dividing the pooled Bayes factor by the Bayes factor from Kerr et al. (2026):

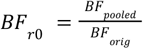

Assuming the pre-registered hypothesis test was examining the presence of an effect, values of *BF*_*r*0_ > 1 indicated replication data were more likely to have occurred under *H*_*r*_ than *H*_0_, indicating a successful replication. Values of *BF*_*r*0_ < 1 indicate the inverse, that the effect has disappeared, therefore an unsuccessful replication. General convention would indicate that values above 3 or below ⅓ indicate a substantial effect, akin to a p-value significance (Stefan et al., 2019). If the pre-registered hypothesis test was confirming no effect existed, these interpretations would reverse, where *BF*_*r*0_ > 1 would indicate a failed replication.

### Heritability

#### Univariate

The best fitting model was re-run on the full sample of twin pairs, and median point estimates of individual level learning rate parameters calculated for all matched twins. All variables were standardised using the scales package (version 1.4.0), before intraclass correlations were calculated for MZ and DZ twins for each learning rate parameter estimate, using the function ICC from the psych package (version 2.5.6). Confidence intervals were estimated using bootstrapping, where twin pairs were resampled with replacement 10,000 times, and the 2.5th and 97.5th percentiles of the bootstrapped distribution used as the 95% intervals.

Formal ACE twin modelling was applied to decompose each learning rate phenotype into latent factors. Each learning rate parameter estimate, p, for each twin, j, in each family, i, was modelled as:

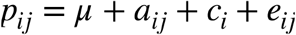

Where μ represented the population mean, *a*_*ij*_ represented additive genetic influences, *c*_*i*_ represented shared environmental influences, and *e*_*ij*_ represented non-shared environmental influences and error, all specific to each twin. Each component was assumed independent and normally distributed with mean 0.

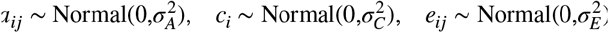

Across all twins, the total phenotypic variance could be decomposed into the separate variances of the three factors, *A, C*, and *E*:

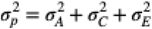

For a vector of two values within a twin pair:

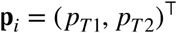

The expected 2 × 2 variance-covariance matrices stored phenotypic variance on the diagonal, and cross-twin covariance on the off-diagonal. It was assumed that *C* was identical within twin pairs, that DZ twins shared approximately half their segregating alleles whereas MZ twins shared all alleles, and that *E* was unique in each twin.

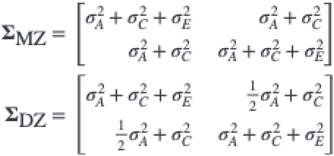

Linear structural equation models (SEM) implemented through standard scripts in openMX (version 2.22.10) estimated these factors through full information maximum likelihood (i.e. finding the values of *A, C*, and *E* which best fit the expected variance-covariance matrices) (Bates et al., 2019). Confidence intervals were calculated using bootstrapping, rather than the likelihood-based method within openMX. For each model, twins pairs were resampled with replacement 1,000 times, with each model refitted to each iterative resampled dataset. Confidence intervals were defined as the 2.5th and 97.5th percentiles of the parameter estimates. Nested submodels, dropping one or more factors (i.e. factors are fixed as 0) (AE, dropping *C*; CE dropping *A*; and E, dropping *A* and *C*) were subsequently fitted, and compared to the full ACE model via likelihood ratio testing:

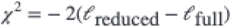

Where *l* represented the summed log-likelihood of the model. Differences in degrees of freedom were equal to the difference in the number of dropped factors. If dropping a factor significantly reduced the fit (i.e. **Δ***χ*^2^ was significant), the more complex model was retained, otherwise the more parsimonious model was chosen (Rijsdijk & Sham, 2002). In a hierarchical manner, the ACE model was first compared to the AE and CE models, before the AE and CE models were compared to the E model. Where two models were tied, i.e. AE and CE both equally well fitting, the model with the lower AIC value was preferred. In the event of tied AIC, theoretical considerations, such as the MZ and DZ ICC values, were used to decide the model selected.

Finally, the factor estimates from the winning model were used to calculate the standardised proportions of total phenotypic variance:

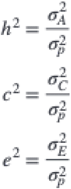

#### Bivariate

To test for shared variance between GAD-7 anxiety severity and appropriate learning parameters, cross-twin cross-trait Pearson correlations were calculated (the correlation between GAD-7 in twin A, and the learning rate parameter in twin B, and vice versa). For those learning parameters showing a significant phenotypic correlation, and a significant cross-twin cross-trait correlation with GAD-7 anxiety severity, bivariate ACE modelling was applied. This extended the approach used in the univariate modelling, now decomposing the covariance between traits into contributions from shared genetic and environmental factors, and estimating the genetic and environmental correlations between traits.

For each individual twin, a parameter vector of the two traits (threat extinction learning rate, and GAD-7 anxiety severity) was created.

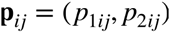

This vector is assumed to decompose into a population mean and three latent sources of variation.

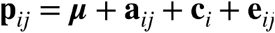

Where each component was assumed to be independent, and follow a multivariate normal distribution.

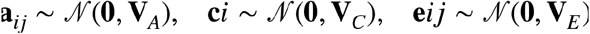

Given the independence of components, the covariance between traits within an individual decomposes analogously.

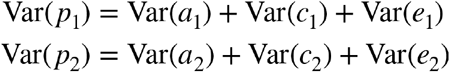

Similarly, the covariance between threat learning rate and GAD-7 anxiety severity within an individual was assumed to decompose similarly.

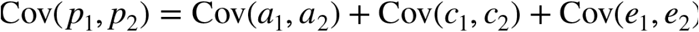

This expands the univariate model by additionally estimating the covariance between genetic (and environmental) influences on the two traits, quantifying the degree to which the same latent factors contribute to both. A 2 × 2 symmetric variance-covariance matrix for each source was created, for example for additive genetic influences:

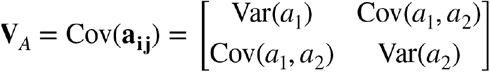

Where the diagonal stored the genetic variance of either trait, and the off diagonal stored the genetic covariance between traits. For a twin pair, with four variables:

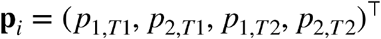

The expected 4 × 4 population covariance matrix (an expansion of the 2 × 2 univariate model matrix) was constructed with the within twin matrices on the diagonal, and the cross-twin matrices on the off diagonal. As in the univariate model, the genetic contribution is halved in the DZ matrix, to facilitate the separation of genetic and environmental influences.

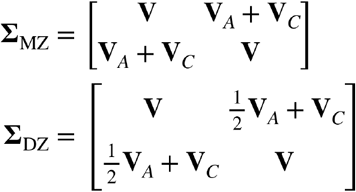

Finally, the cross-trait genetic and environmental correlations were calculated by standardising the off-diagonal element by the standard deviations of each trait, for example the genetic correlation:

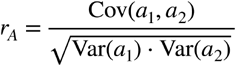

This quantified the extent to which the same genetic factors influence both GAD-7 anxiety severity and threat extinction learning rate, with a higher value indicating greater genetic overlap. As in the univariate approach, nested submodels were created by constraining the entire 2 × 2 variance-covariance matrices to 0, i.e. the AE model:

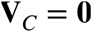

Each submodel was compared to the full ACE model identically to the univariate approach.

#### Hypotheses

Given the novelty of our approach, and lack of prior empirical data, univariate predictions were simply of overall effect size, rather than proposing comparisons between phases and CS.

*H*_*4*_: Threat positive learning rate, threat negative learning rate, threat extinction learning rate, safety learning rate, and safety confirmation learning rate will display moderate univariate heritability (*h*^2^) of between 0.3 and 0.5.

*H*_*5*_: Threat negative learning rate, threat extinction learning rate, safety learning rate will display moderate bivariate genetic correlations, *r*_*A*_, of between 0.3 and 0.5, with GAD-7 anxiety severity.

## Declarations

O.J.R has completed consultancy work for Peak, IESO digital health, Roche and BlackThorn therapeutics and sat on the committee of the British Association for Psychopharmacology until 2022.

## Acknowledgements

O.J.R is funded by an ERC Advanced Grant (UKRI underwrite).

T.C.E is part-funded by the National Institute for Health and Care Research (NIHR) Maudsley Biomedical Research Centre (BRC). The views expressed are those of the author(s) and not necessarily those of the NHS, the NIHR or the Department of Health.

For the purposes of open access, the author has applied a Creative Commons Attribution (CC BY) licence to any Accepted Author Manuscript version arising from this submission

## Data availability

The data used in this study are not publicly available per the terms and conditions of the Twins Early Development Study. However, data is available upon request and subsequent acceptance of a data sharing agreement. Requests should be made via https://datarequests.teds.ac.uk/.

## Code availability

All code is available on this study’s OSF page https://osf.io/jsmak/.

## Notes

https://osf.io/jsmak/overview

