## Supplemental tables for "The heritability of reinforcement learning parameters and their association with anxiety"

### Supplementary tables

#### Supplementary table 1. Task exclusion criteria.

| Criteria | Strict | Moderate |
| --- | --- | --- |
| n | 1151 | 1357 |
| Complete pairs (MZ;DZ) | 108;118 | 138;160 |
| Did not complete extinction phase | ✓ | ✓ |
| Removed headphones | ✓ | ✓ |
| Restarted app after trial one | ✓ |  |
| Rated US unpleasantness <= 5 | ✓ | ✓ |
| Contingency unaware | ✓ |  |
| Volume below 100% (any trial) | ✓ |  |
| Volume below 80% (any trial) | ✓ | ✓ |
| Stated non-compliance with instructions | ✓ | ✓ |

#### Supplementary table 2. Overall model comparison.

| **Model** | **Learning rate configuration** | **LR count** | **Subject params** | **Total params** | **Group params** | **Log likelihood** | **LOOIC** | **WAIC** | **Pseudo R²** | **ELPD LOO** |
| --- | --- | --- | --- | --- | --- | --- | --- | --- | --- | --- |
| 1 | Single | 1 | 5 | 4,641 | 16 | -61,782.8 | 133,535.8 | 130,578.8 | 0.4876 | -66,767.9 |
| 2 | CS+, CS- | 2 | 6 | 5,568 | 18 | -60,559.5 | 131,261.9 | 128,070.1 | 0.4978 | -65,631.0 |
| 3 | CS+ Acq., CS+ Ext., CS- | 3 | 7 | 6,495 | 20 | -58,994.2 | 128,339.8 | 125,019.4 | 0.5108 | -64,169.9 |
| 4 | CS+ Acq. CS+ Ext., CS- Acq., CS- Ext. | 4 | 8 | 7,422 | 22 | -58,129.7 | 126,846.8 | 123,451.8 | 0.5179 | -63,423.4 |
| 5 | US+, US- | 2 | 6 | 5,568 | 18 | -59,223.1 | 130,222.4 | 126,655.4 | 0.5089 | -65,111.2 |
| 6 | CS+US+, CS+US-, CS- | 3 | 7 | 6,495 | 20 | -57,583.2 | 126,613.6 | 122,966.4 | 0.5225 | -63,306.8 |
| 7 | CS+US+ Acq., CS+US- Acq., CS+ Ext. CS- Acq., CS- Ext. | 5 | 9 | 8,349 | 24 | **-55,546.5** | **122,908.0** | **119,048.0** | **0.5394** | **-61,454.0** |

• ELPD-LOO: Expected Log Predictive Density; LOOIC: Leave-One-Out Information Criterion; SE: Standard error of LOOIC estimate; WAIC: Widely Applicable Information Criterion; Pseudo R²: variance explained vs chance; LL: log likelihood; LOO: Leave-One-Out cross-validation estimate

• Bold text = overall best model

#### Supplementary table 3. Model comparison for each phase-CS combination.

| **Model** | **Learning Rate Configuration** | **Log Likelihood** | **LOOIC** | **WAIC** | **Pseudo R²** | **ELPD LOO** |
| --- | --- | --- | --- | --- | --- | --- |
|  | ***Acquisition CS+*** |  |  |  |  |  |
| 1 | Single | -15549.4 | 33574.5 | 32847.1 | 0.3532 | -16787.3 |
| 2 | CS+, CS- | -15267.5 | 33047.1 | 32265.7 | 0.3649 | -16523.6 |
| 3 | CS+ Acq., CS+ Ext., CS- | -14910 | 32446 | 31620.1 | 0.3798 | -16223 |
| 4 | CS+ Acq. CS+ Ext., CS- Acq., CS- Ext. | -14690.8 | 32067.9 | 31201.5 | 0.3889 | -16033.9 |
| 5 | US+, US- | -14926.2 | 32848.8 | 31963.7 | 0.3791 | -16424.4 |
| 6 | CS+US+, CS+US-, CS- | -14550.3 | 31954.2 | 31075.3 | 0.3947 | -15977.1 |
| 7 | CS+US+ Acq., CS+US- Acq., CS+ Ext. CS- Acq., CS- Ext. | **-14029.8** | **31043** | **30100.2** | **0.4164** | **-15521.5** |
|  | ***Acquisition CS-*** |  |  |  |  |  |
| 1 | Single | -15470.9 | 33486.2 | 32705.5 | 0.3537 | -16743.1 |
| 2 | CS+, CS- | -15145.7 | 32835.8 | 32024.8 | 0.3673 | -16417.9 |
| 3 | CS+ Acq., CS+ Ext., CS- | -14739 | 32066.4 | 31214.2 | 0.3843 | -16033.2 |
| 4 | CS+ Acq. CS+ Ext., CS- Acq., CS- Ext. | -14545.8 | 31726.2 | 30887 | 0.3924 | -15863.1 |
| 5 | US+, US- | -14814.9 | 32557.9 | 31645.9 | 0.3811 | -16279 |
| 6 | CS+US+, CS+US-, CS- | -14386.2 | 31609.7 | 30682.3 | 0.399 | -15804.9 |
| 7 | CS+US+ Acq., CS+US- Acq., CS+ Ext. CS- Acq., CS- Ext. | **-13892.9** | **30738.1** | **29755.8** | **0.4196** | **-15369.1** |
|  | ***Extinction CS+*** |  |  |  |  |  |
| 1 | Single | -15499 | 33470 | 32761.6 | 0.5732 | -16735 |
| 2 | CS+, CS- | -15134.4 | 32809 | 32010.4 | 0.5832 | -16404.5 |
| 3 | CS+ Acq., CS+ Ext., CS- | -14704.5 | 31950.5 | 31142.1 | 0.595 | -15975.2 |
| 4 | CS+ Acq. CS+ Ext., CS- Acq., CS- Ext. | -14478.5 | 31553.8 | 30750.1 | 0.6013 | -15776.9 |
| 5 | US+, US- | -14838.5 | 32608 | 31714.8 | 0.5914 | -16304 |
| 6 | CS+US+, CS+US-, CS- | -14380.8 | 31629.1 | 30716.9 | 0.604 | -15814.5 |
| 7 | CS+US+ Acq., CS+US- Acq., CS+ Ext. CS- Acq., CS- Ext. | **-13888.7** | **30702.1** | **29756.6** | **0.6175** | **-15351** |
|  | ***Extinction CS-*** |  |  |  |  |  |
| 1 | Single | -15263.5 | 32996 | 32264.5 | 0.5794 | -16498 |
| 2 | CS+, CS- | -15011.8 | 32570.9 | 31769.2 | 0.5864 | -16285.5 |
| 3 | CS+ Acq., CS+ Ext., CS- | -14640.7 | 31875.3 | 31043 | 0.5966 | -15937.6 |
| 4 | CS+ Acq. CS+ Ext., CS- Acq., CS- Ext. | -14414.5 | 31499.9 | 30613.1 | 0.6028 | -15750 |
| 5 | US+, US- | -14643.5 | 32201.2 | 31331 | 0.5965 | -16100.6 |
| 6 | CS+US+, CS+US-, CS- | -14265.9 | 31414.2 | 30492 | 0.6069 | -15707.1 |
| 7 | CS+US+ Acq., CS+US- Acq., CS+ Ext. CS- Acq., CS- Ext. | **-13735.2** | **30424.6** | **29435.4** | **0.6216** | **-15212.3** |

• ELPD-LOO: Expected Log Predictive Density; LOOIC: Leave-One-Out Information Criterion; SE: Standard error of LOOIC estimate; WAIC: Widely Applicable Information Criterion; Pseudo R²: variance explained vs chance; LL: log likelihood; LOO: Leave-One-Out cross-validation estimate

• Bold text = overall best model

#### Supplementary table 4. Real and model generated descriptive measure correlation with GAD-7.

|  | Participant data (strict) | | Model generated data (strict) | | Participant data (moderate) | | Model generated data (moderate) | |
| --- | --- | --- | --- | --- | --- | --- | --- | --- |
| Whole phase mean | ⍴GAD-7 [95% CI] | p | ρGAD-7 [95% CI] | p | ρGAD-7 [95% CI] | p | ρGAD-7 [95% CI] | p |
| Acquisition CS+ | -0.01 [-0.08, 0.06] | .816 | 0.04 [-0.03, 0.11] | .211 | -0.06 [-0.12, 0.00] | .056 | -0.01 [-0.07, 0.06] | .843 |
| Acquisition CS- | **0.07 [0.01, 0.14]** | .025 | 0.06 [-0.01, 0.13] | .068 | **0.11 [0.05, 0.17]** | **< .001** | **0.10 [0.03, 0.16]** | **.002** |
| Extinction CS+ | **0.13 [0.07, 0.20]** | < .001 | **0.13 [0.07, 0.20]** | **< .001** | **0.13 [0.07, 0.19]** | **< .001** | **0.13 [0.07, 0.19]** | **< .001** |
| Extinction CS- | 0.05 [-0.02, 0.12] | .124 | 0.05 [-0.01, 0.12] | .128 | 0.05 [-0.01, 0.11] | .098 | **0.06 [-0.00, 0.12]** | **.044** |

Spearman correlation between GAD-7 anxiety severity, and descriptive fear conditioning measures (mean of whole phase responding per CS). Correlations that exist within participant responding should be reproduced by model generated data, if the model is a valid analogue of the data generating process. In the strict exclusion group, the small correlation noted in safety learning is not quite reproduced by the model, indicating a less than ideal fit. This is not the case in the moderate exclusion group.

#### Supplementary table 5. Correlation and residual analysis (strict exclusion criteria).

|  | GAD-7 | | PHQ-8 | | GAD-7 Unique | | PHQ-8 Unique | | Shared | |
| --- | --- | --- | --- | --- | --- | --- | --- | --- | --- | --- |
| Parameter | ρ [95% CI] | p | ρ [95% CI] | p | ρ [95% CI] | p | ρ [95% CI] | p | ρ [95% CI] | p |
| Threat positive | -0.06 [-0.13, 0.00] | 0.057 | -0.03 [-0.09, 0.04] | 0.418 | -0.07 [-0.13, -0.00] | 0.034 | 0.04 [-0.03, 0.10] | 0.270 | -0.04 [-0.11, 0.02] | 0.172 |
| Threat negative | -0.08 [-0.15, -0.02] | 0.011 | -0.06 [-0.13, 0.00] | 0.052 | -0.05 [-0.12, 0.02] | 0.131 | 0.00 [-0.06, 0.07] | 0.952 | -0.08 [-0.14, -0.01] | 0.019 |
| Safety | -0.07 [-0.13, -0.00] | 0.045 | -0.08 [-0.15, -0.02] | 0.011 | -0.00 [-0.06, 0.06] | 0.980 | -0.06 [-0.12, 0.01] | 0.088 | -0.08 [-0.14, -0.01] | 0.015 |
| Threat extinction | **-0.14 [-0.21, -0.08]** | **0.000** | **-0.14 [-0.21, -0.07]** | **0.000** | -0.06 [-0.13, 0.00] | 0.056 | -0.06 [-0.13, 0.00] | 0.049 | **-0.15 [-0.22, -0.08]** | **0.000** |
| Safety confirmation | **-0.12 [-0.18, -0.05]** | **0.000** | **-0.10 [-0.17, -0.04]** | **0.002** | -0.06 [-0.12, 0.01] | 0.084 | -0.04 [-0.10, 0.03] | 0.261 | **-0.12 [-0.18, -0.05]** | **0.000** |
| CS+ Initial Value | 0.05 [-0.01, 0.12] | 0.112 | 0.02 [-0.04, 0.09] | 0.477 | 0.05 [-0.01, 0.12] | 0.111 | -0.03 [-0.10, 0.03] | 0.326 | 0.04 [-0.02, 0.10] | 0.238 |
| CS– Initial Value | 0.05 [-0.01, 0.12] | 0.110 | 0.01 [-0.05, 0.08] | 0.751 | **0.09 [0.03, 0.16]** | **0.005** | -0.05 [-0.12, 0.01] | 0.103 | 0.03 [-0.03, 0.09] | 0.350 |
| Extinction CS– Jump | -0.05 [-0.12, 0.01] | 0.102 | -0.04 [-0.11, 0.02] | 0.178 | -0.03 [-0.09, 0.04] | 0.389 | -0.02 [-0.08, 0.05] | 0.641 | -0.05 [-0.12, 0.01] | 0.110 |
| Response precision (φ) | -0.05 [-0.11, 0.01] | 0.129 | -0.03 [-0.10, 0.03] | 0.319 | -0.04 [-0.11, 0.02] | 0.182 | 0.02 [-0.04, 0.08] | 0.566 | -0.04 [-0.11, 0.02] | 0.206 |
| *Bold indicates p < 0.008 (Bonferroni-corrected α = 0.05/6)* | | | | | | | | | | |
| *Values show Spearman ρ [95% bootstrap CI]* | | | | | | | | | | |
| *N = 925 (925 group)* | | | | | | | | | | |
| *GAD-7 = Generalised Anxiety Disorder 7-item; PHQ-8 = Patient Health Questionnaire 8-item* | | | | | | | | | | |
| *Unique = variance after partialling out the other measure; Shared = common variance (mean of z-scored ranks)* | | | | | | | | | | |

#### Supplementary table 6. Correlation and residual analysis (moderate exclusion criteria).

|  | GAD-7 | | PHQ-8 | | GAD-7 Unique | | PHQ-8 Unique | | Shared | |
| --- | --- | --- | --- | --- | --- | --- | --- | --- | --- | --- |
| Parameter | ρ [95% CI] | p | ρ [95% CI] | p | ρ [95% CI] | p | ρ [95% CI] | p | ρ [95% CI] | p |
| Threat positive | **-0.11 [-0.17, -0.05]** | **0.000** | -0.07 [-0.13, -0.01] | 0.026 | **-0.10 [-0.16, -0.04]** | **0.001** | 0.03 [-0.03, 0.09] | 0.354 | **-0.09 [-0.15, -0.03]** | **0.003** |
| Threat negative | **-0.09 [-0.15, -0.03]** | **0.003** | -0.06 [-0.12, 0.00] | 0.055 | -0.07 [-0.14, -0.01] | 0.016 | 0.03 [-0.04, 0.09] | 0.402 | -0.08 [-0.14, -0.02] | 0.010 |
| Safety | **-0.09 [-0.15, -0.03]** | **0.002** | **-0.08 [-0.14, -0.02]** | **0.007** | -0.04 [-0.10, 0.02] | 0.156 | -0.03 [-0.09, 0.03] | 0.395 | **-0.09 [-0.15, -0.03]** | **0.002** |
| Threat extinction | **-0.16 [-0.22, -0.10]** | **0.000** | **-0.13 [-0.19, -0.07]** | **0.000** | **-0.10 [-0.16, -0.04]** | **0.001** | -0.03 [-0.08, 0.03] | 0.411 | **-0.15 [-0.21, -0.09]** | **0.000** |
| Safety confirmation | **-0.10 [-0.17, -0.04]** | **0.001** | **-0.10 [-0.16, -0.04]** | **0.001** | -0.04 [-0.10, 0.02] | 0.232 | -0.05 [-0.10, 0.01] | 0.137 | **-0.11 [-0.17, -0.05]** | **0.000** |
| CS+ Initial Value | 0.04 [-0.02, 0.10] | 0.226 | 0.02 [-0.04, 0.08] | 0.593 | 0.04 [-0.03, 0.10] | 0.240 | -0.03 [-0.09, 0.03] | 0.368 | 0.03 [-0.03, 0.09] | 0.354 |
| CS– Initial Value | **0.11 [0.05, 0.17]** | **0.000** | 0.06 [-0.00, 0.12] | 0.065 | **0.13 [0.07, 0.19]** | **0.000** | -0.05 [-0.11, 0.01] | 0.084 | **0.09 [0.02, 0.15]** | **0.005** |
| Extinction CS– Jump | -0.07 [-0.13, -0.01] | 0.030 | -0.05 [-0.11, 0.01] | 0.132 | -0.05 [-0.11, 0.01] | 0.092 | 0.00 [-0.06, 0.06] | 0.949 | -0.06 [-0.12, 0.00] | 0.052 |
| Response precision (φ) | **-0.09 [-0.15, -0.03]** | **0.002** | -0.07 [-0.14, -0.01] | 0.015 | -0.06 [-0.12, -0.00] | 0.043 | 0.00 [-0.06, 0.06] | 0.946 | **-0.09 [-0.15, -0.02]** | **0.005** |
| *Bold indicates p < 0.008 (Bonferroni-corrected α = 0.05/6)* | | | | | | | | | | |
| *Values show Spearman ρ [95% bootstrap CI]* | | | | | | | | | | |
| *N = 1059 (1059 group)* | | | | | | | | | | |
| *GAD-7 = Generalised Anxiety Disorder 7-item; PHQ-8 = Patient Health Questionnaire 8-item* | | | | | | | | | | |
| *Unique = variance after partialling out the other measure; Shared = common variance (mean of z-scored ranks)* | | | | | | | | | | |

#### Supplementary table 7. Bayes replication factor analysis (strict exclusion criteria).

| Measure | Parameter | n orig. | n rep | BF₁₀ | BF pooled | BF r0 | Posterior ρ [95% HDI] | Interpretation | Hypothesis | H_1_ favoured? |
| --- | --- | --- | --- | --- | --- | --- | --- | --- | --- | --- |
| GAD-7 | Threat negative | 88 | 925 | **10.69** | 13.93 | 1.30 | -0.10 [-0.16, -0.04] | Inconclusive | H2c, ρ<0 | ╳ |
|  | Safety | 88 | 925 | **17.53** | 3.51 | 0.20 | -0.09 [-0.15, -0.03] | Effect disappeared | H2a, ρ<0 | ╳ |
|  | Threat extinction | 88 | 925 | **28.93** | 34413.07 | **1189.67** | -0.16 [-0.22, -0.10] | Effect replicated | H2b, ρ<0 | ✓ |
| PHQ-8 | Threat negative | 88 | 925 | 1.22 | 1.29 | 1.06 | -0.08 [-0.14, -0.01] | Inconclusive | H2f, ρ=0 | ✓ |
|  | Safety | 88 | 925 | 0.54 | 3.78 | **6.95** | -0.09 [-0.15, -0.03] | Effect emerged | H2d, ρ=0 | ╳ |
|  | Threat extinction | 88 | 925 | **3.70** | 7184.97 | **1939.58** | -0.15 [-0.21, -0.09] | Effect replicated | H2e, ρ<0 | ✓ |
| *Strict exclusion: pilot N=88, replication N=925*  *BF₁₀ = original; BF r0 = replication (pooled/original)* | | | | | | | | | | |

#### Supplementary table 8. Bayes replication factor analysis (moderate exclusion criteria).

| Measure | Parameter | n orig. | n rep | BF₁₀ | BF pooled | BF r0 | Posterior ρ [95% HDI] | Interpretation | Hypothesis | H_1_ favoured? |
| --- | --- | --- | --- | --- | --- | --- | --- | --- | --- | --- |
| GAD-7 | Threat negative | 145 | 1,059 | 0.37 | 11.28 | **30.77** | -0.09 [-0.15, -0.04] | Effect emerged | H3c, ρ=0 | ╳ |
|  | Safety | 145 | 1,059 | **5.44** | 74.90 | **13.76** | -0.11 [-0.16, -0.05] | Effect replicated | H3a, ρ<ρ_strict_ | ╳ |
|  | Threat extinction | 145 | 1,059 | **4.76** | 1350649.01 | **283655.07** | -0.17 [-0.22, -0.11] | Effect replicated | H3b, ρ<ρ_strict_ | ╳ |
| PHQ-8 | Threat negative | 145 | 1,059 | 0.24 | 0.54 | 2.24 | -0.06 [-0.11, -0.00] | Inconclusive | H3f, ρ=0 | ✓ |
|  | Safety | 145 | 1,059 | 0.70 | 7.94 | **11.36** | -0.09 [-0.15, -0.03] | Effect emerged | H3d, ρ=0 | ╳ |
|  | Threat extinction | 145 | 1,059 | **7.00** | 13015.97 | **1858.67** | -0.14 [-0.20, -0.09] | Effect replicated | H3e, ρ<ρ_strict_ | ✓ |
| *Moderate exclusion: pilot N=145, replication N=1059*  *BF₁₀ = original; BF r0 = replication (pooled/original)* | | | | | | | | | | |

#### Supplementary table 9. Twin demographics.

| **Exclusion Criteria** | **n** | **Female n (%)** | **Age Mean (SD)** | **GAD-7 Median (IQR)** | **PHQ-8 Median (IQR)** |
| --- | --- | --- | --- | --- | --- |
| Strict Exclusion | 452 | 322 (71.2%) | 23.7 (0.9) | 4 (5) | 4 (6) |
| Moderate Exclusion | 596 | 432 (72.5%) | 23.7 (0.9) | 4 (5) | 4 (6) |
| *Comparison (Strict vs Moderate)* | n = 452 / 596 / | χ²=0.14, p=0.708 | W=132461, p=0.645 | W=137221, p=0.601 | W=134752, p=0.991 |
| *Bold indicates p < 0.05* | | | | | |
| *Kruskal-Wallis test used for continuous variables; Chi-square test used for categorical variables* | | | | | |
| *GAD-7 = Generalised Anxiety Disorder 7-item scale; PHQ-8 = Patient Health Questionnaire 9-item scale* | | | | | |

#### Supplementary table 10. Twin correlations, ACE model fitting, and variance estimates (moderate exclusion criteria).

| **MZ ICC (n=138)** | **DZ ICC (n=160)** | **Model** | **A** | **C** | **E** | **-2LL** | **df** | **AIC** | **BIC** | **χ² vs ACE** | **p vs ACE** | **χ² vs AE** | **p vs AE** | **χ² vs CE** | **p vs CE** |
| --- | --- | --- | --- | --- | --- | --- | --- | --- | --- | --- | --- | --- | --- | --- | --- |
| **Threat positive** | | | | | | | | | | | | | | | |
| **0.18 (0.03, 0.31)** | 0.05 (-0.10, 0.20) | ACE | 0.26 (-0.14, 0.66) | -0.08 (-0.41, 0.23) | 0.82 (0.68, 0.97) | 1685.82 | 592 | 501.82 | -1686.86 | - | - | - | - | - | - |
|  |  | **AE** | **0.16 (0.01, 0.28)** | **0.00 (0.00, 0.00)** | **0.84 (0.72, 0.99)** | **1686.03** | **593** | **500.03** | **-1692.34** | **0.22** | **0.642** | **-** | **-** | **-** | **-** |
|  |  | CE | 0.00 (0.00, 0.00) | 0.10 (-0.00, 0.21) | 0.90 (0.79, 1.00) | 1687.09 | 593 | 501.09 | -1691.29 | 1.27 | 0.260 | - | - | - | - |
|  |  | E | 0.00 (0.00, 0.00) | 0.00 (0.00, 0.00) | 1.00 (1.00, 1.00) | 1690.37 | 594 | 502.37 | -1693.70 | 4.56 | 0.102 | *4.34* | *0.037* | 3.29 | 0.070 |
| **Threat negative** | | | | | | | | | | | | | | | |
| 0.00 (-0.17, 0.20) | 0.08 (-0.12, 0.27) | ACE | -0.17 (-0.73, 0.42) | 0.17 (-0.32, 0.65) | 1.00 (0.79, 1.16) | 1689.42 | 592 | 505.42 | -1683.26 | - | - | - | - | - | - |
|  |  | AE | 0.03 (-0.12, 0.20) | 0.00 (0.00, 0.00) | 0.97 (0.80, 1.12) | 1690.22 | 593 | 504.22 | -1688.16 | 0.80 | 0.370 | - | - | - | - |
|  |  | CE | 0.00 (0.00, 0.00) | 0.04 (-0.10, 0.18) | 0.96 (0.82, 1.10) | 1689.95 | 593 | 503.95 | -1688.43 | 0.53 | 0.465 | - | - | - | - |
|  |  | **E** | **0.00 (0.00, 0.00)** | **0.00 (0.00, 0.00)** | **1.00 (1.00, 1.00)** | **1690.37** | **594** | **502.37** | **-1693.70** | **0.96** | **0.619** | **0.15** | **0.694** | **0.42** | **0.515** |
| **Safety** | | | | | | | | | | | | | | | |
| **0.25 (0.08, 0.40)** | 0.04 (-0.12, 0.20) | ACE | 0.39 (-0.08, 0.81) | -0.16 (-0.51, 0.23) | 0.77 (0.62, 0.92) | 1681.56 | 592 | 497.56 | -1691.12 | - | - | - | - | - | - |
|  |  | **AE** | **0.21 (0.07, 0.34)** | **0.00 (0.00, 0.00)** | **0.79 (0.66, 0.93)** | **1682.30** | **593** | **496.30** | **-1696.08** | **0.74** | **0.388** | **-** | **-** | **-** | **-** |
|  |  | CE | 0.00 (0.00, 0.00) | 0.14 (0.03, 0.25) | 0.86 (0.75, 0.97) | 1684.56 | 593 | 498.56 | -1693.82 | 3.00 | 0.083 | - | - | - | - |
|  |  | E | 0.00 (0.00, 0.00) | 0.00 (0.00, 0.00) | 1.00 (1.00, 1.00) | 1690.37 | 594 | 502.37 | -1693.70 | *8.82* | *0.012* | *8.07* | *0.004* | *5.81* | *0.016* |
| **Threat extinction** | | | | | | | | | | | | | | | |
| 0.14 (-0.02, 0.27) | 0.12 (-0.03, 0.27) | ACE | 0.08 (-0.36, 0.48) | 0.07 (-0.24, 0.39) | 0.85 (0.70, 1.02) | 1685.47 | 592 | 501.47 | -1687.21 | - | - | - | - | - | - |
|  |  | AE | 0.17 (0.02, 0.31) | 0.00 (0.00, 0.00) | 0.83 (0.69, 0.98) | 1685.65 | 593 | 499.65 | -1692.73 | 0.18 | 0.674 | - | - | - | - |
|  |  | **CE** | **0.00 (0.00, 0.00)** | **0.13 (0.02, 0.22)** | **0.87 (0.78, 0.98)** | **1685.59** | **593** | **499.59** | **-1692.79** | **0.12** | **0.732** | **-** | **-** | **-** | **-** |
|  |  | E | 0.00 (0.00, 0.00) | 0.00 (0.00, 0.00) | 1.00 (1.00, 1.00) | 1690.37 | 594 | 502.37 | -1693.70 | 4.90 | 0.086 | *4.73* | *0.030* | *4.79* | *0.029* |
| **Safety confirmation** | | | | | | | | | | | | | | | |
| **0.19 (0.02, 0.33)** | 0.06 (-0.09, 0.21) | ACE | 0.28 (-0.18, 0.69) | -0.08 (-0.41, 0.27) | 0.81 (0.65, 0.98) | 1684.98 | 592 | 500.98 | -1687.70 | - | - | - | - | - | - |
|  |  | **AE** | **0.17 (0.03, 0.32)** | **0.00 (0.00, 0.00)** | **0.83 (0.68, 0.97)** | **1685.20** | **593** | **499.20** | **-1693.18** | **0.22** | **0.638** | **-** | **-** | **-** | **-** |
|  |  | CE | 0.00 (0.00, 0.00) | 0.11 (0.00, 0.22) | 0.89 (0.78, 1.00) | 1686.43 | 593 | 500.43 | -1691.95 | 1.45 | 0.228 | - | - | - | - |
|  |  | E | 0.00 (0.00, 0.00) | 0.00 (0.00, 0.00) | 1.00 (1.00, 1.00) | 1690.37 | 594 | 502.37 | -1693.70 | 5.40 | 0.067 | *5.18* | *0.023* | *3.95* | *0.047* |
| **CS+ Initial Value** | | | | | | | | | | | | | | | |
| 0.01 (-0.17, 0.18) | 0.11 (-0.05, 0.26) | ACE | -0.21 (-0.64, 0.24) | 0.21 (-0.14, 0.54) | 1.00 (0.83, 1.15) | 1688.57 | 592 | 504.57 | -1684.11 | - | - | - | - | - | - |
|  |  | AE | 0.05 (-0.10, 0.19) | 0.00 (0.00, 0.00) | 0.95 (0.81, 1.10) | 1689.96 | 593 | 503.96 | -1688.42 | 1.39 | 0.238 | - | - | - | - |
|  |  | CE | 0.00 (0.00, 0.00) | 0.06 (-0.05, 0.17) | 0.94 (0.83, 1.05) | 1689.40 | 593 | 503.40 | -1688.97 | 0.84 | 0.361 | - | - | - | - |
|  |  | **E** | **0.00 (0.00, 0.00)** | **0.00 (0.00, 0.00)** | **1.00 (1.00, 1.00)** | **1690.37** | **594** | **502.37** | **-1693.70** | **1.81** | **0.405** | **0.41** | **0.520** | **0.97** | **0.324** |
| **CS- Initial Value** | | | | | | | | | | | | | | | |
| -0.08 (-0.25, 0.08) | 0.03 (-0.13, 0.19) | ACE | -0.22 (-0.72, 0.23) | 0.14 (-0.21, 0.53) | 1.08 (0.93, 1.24) | 1689.27 | 592 | 505.27 | -1683.41 | - | - | - | - | - | - |
|  |  | AE | -0.05 (-0.19, 0.09) | 0.00 (0.00, 0.00) | 1.05 (0.91, 1.19) | 1689.85 | 593 | 503.85 | -1688.52 | 0.59 | 0.444 | - | - | - | - |
|  |  | CE | 0.00 (0.00, 0.00) | -0.03 (-0.14, 0.09) | 1.03 (0.91, 1.14) | 1690.18 | 593 | 504.18 | -1688.20 | 0.91 | 0.339 | - | - | - | - |
|  |  | **E** | **0.00 (0.00, 0.00)** | **0.00 (0.00, 0.00)** | **1.00 (1.00, 1.00)** | **1690.37** | **594** | **502.37** | **-1693.70** | **1.11** | **0.575** | **0.52** | **0.470** | **0.20** | **0.658** |
| **Extinction CS- Jump** | | | | | | | | | | | | | | | |
| **0.28 (0.10, 0.43)** | 0.06 (-0.11, 0.21) | ACE | 0.43 (0.00, 0.86) | -0.16 (-0.51, 0.18) | 0.73 (0.57, 0.88) | 1679.12 | 592 | 495.12 | -1693.56 | - | - | - | - | - | - |
|  |  | **AE** | **0.24 (0.09, 0.40)** | **0.00 (0.00, 0.00)** | **0.76 (0.60, 0.91)** | **1679.96** | **593** | **493.96** | **-1698.41** | **0.84** | **0.359** | **-** | **-** | **-** | **-** |
|  |  | CE | 0.00 (0.00, 0.00) | 0.16 (0.05, 0.27) | 0.84 (0.73, 0.95) | 1682.90 | 593 | 496.90 | -1695.47 | 3.78 | 0.052 | - | - | - | - |
|  |  | E | 0.00 (0.00, 0.00) | 0.00 (0.00, 0.00) | 1.00 (1.00, 1.00) | 1690.37 | 594 | 502.37 | -1693.70 | *11.25* | *0.004* | *10.41* | *0.001* | *7.47* | *0.006* |
| **Response precision** | | | | | | | | | | | | | | | |
| 0.11 (-0.00, 0.26) | **0.33 (0.11, 0.54)** | ACE | -0.30 (-0.85, 0.46) | 0.44 (-0.04, 0.91) | 0.86 (0.51, 1.00) | 1669.93 | 592 | 485.93 | -1702.74 | - | - | - | - | - | - |
|  |  | AE | 0.29 (0.11, 0.52) | 0.00 (0.00, 0.00) | 0.71 (0.48, 0.89) | 1677.25 | 593 | 491.25 | -1701.13 | *7.31* | *0.007* | - | - | - | - |
|  |  | **CE** | **0.00 (0.00, 0.00)** | **0.25 (0.09, 0.40)** | **0.75 (0.60, 0.91)** | **1671.54** | **593** | **485.54** | **-1706.84** | **1.60** | **0.205** | **-** | **-** | **-** | **-** |
|  |  | E | 0.00 (0.00, 0.00) | 0.00 (0.00, 0.00) | 1.00 (1.00, 1.00) | 1690.37 | 594 | 502.37 | -1693.70 | *20.44* | *<.001* | *13.13* | *<.001* | *18.84* | *<.001* |
| *95% confidence intervals via percentile bootstrap (ICC: 10000, ACE: 1000 resamples).* | | | | | | | | | | | | | | | |
| *A = additive genetic; C = shared environment; E = unique environment + error.* | | | | | | | | | | | | | | | |
| *Best-fitting model per parameter in bold (hierarchical likelihood ratio testing).* | | | | | | | | | | | | | | | |
| *Bold ICCs indicate significance (95% CI excludes zero).* | | | | | | | | | | | | | | | |
| *Italic chi2/p indicates significantly worse fit (p < .05).* | | | | | | | | | | | | | | | |

#### Supplementary table 11. Cross-twin cross-trait correlations (strict exclusion criteria).

|  | **Phenotypic** | **Within-trait: GAD-7** | | **Within-trait: Parameter** | | **Cross-twin Cross-trait** | | **Pairs** | |
| --- | --- | --- | --- | --- | --- | --- | --- | --- | --- |
| **Learning Parameter** | **r [95% CI]** | **MZ r [95% CI]** | **DZ r [95% CI]** | **MZ r [95% CI]** | **DZ r [95% CI]** | **MZ r [95% CI]** | **DZ r [95% CI]** | **MZ n** | **DZ n** |
| Threat positive | -0.04 [-0.13, 0.05] | **0.53 [0.36, 0.68]** | **0.23 [0.05, 0.43]** | 0.09 [-0.06, 0.24] | -0.02 [-0.19, 0.17] | 0.03 [-0.11, 0.16] | 0.03 [-0.12, 0.17] | 108 | 118 |
| Threat negative | -0.01 [-0.11, 0.10] | **0.53 [0.36, 0.68]** | **0.23 [0.05, 0.43]** | 0.01 [-0.17, 0.23] | 0.17 [-0.06, 0.37] | 0.12 [-0.01, 0.26] | 0.02 [-0.11, 0.16] | 108 | 118 |
| Safety | -0.01 [-0.10, 0.09] | **0.53 [0.36, 0.67]** | **0.23 [0.05, 0.43]** | **0.19 [0.02, 0.36]** | 0.03 [-0.16, 0.22] | -0.11 [-0.23, 0.02] | -0.06 [-0.16, 0.05] | 108 | 118 |
| Threat extinction | **-0.14 [-0.24, -0.05]** | **0.53 [0.36, 0.68]** | **0.23 [0.06, 0.42]** | 0.07 [-0.09, 0.23] | 0.10 [-0.09, 0.28] | -0.00 [-0.12, 0.12] | -0.08 [-0.19, 0.03] | 108 | 118 |
| Safety confirmation | -0.10 [-0.20, 0.01] | **0.53 [0.36, 0.68]** | **0.23 [0.05, 0.42]** | 0.09 [-0.09, 0.26] | 0.03 [-0.14, 0.20] | -0.06 [-0.20, 0.08] | 0.03 [-0.09, 0.17] | 108 | 118 |
| CS+ Initial Value | 0.06 [-0.03, 0.15] | **0.53 [0.36, 0.68]** | **0.23 [0.05, 0.43]** | 0.00 [-0.19, 0.21] | 0.12 [-0.06, 0.31] | -0.08 [-0.20, 0.05] | 0.05 [-0.08, 0.17] | 108 | 118 |
| CS- Initial Value | -0.02 [-0.11, 0.08] | **0.53 [0.36, 0.68]** | **0.23 [0.05, 0.43]** | 0.04 [-0.15, 0.23] | 0.10 [-0.09, 0.28] | -0.03 [-0.16, 0.10] | -0.04 [-0.17, 0.09] | 108 | 118 |
| Extinction CS- Jump | 0.03 [-0.06, 0.11] | **0.53 [0.35, 0.68]** | **0.23 [0.05, 0.42]** | **0.22 [0.03, 0.40]** | 0.15 [-0.03, 0.33] | 0.05 [-0.09, 0.18] | 0.06 [-0.07, 0.20] | 108 | 118 |
| Response precision | -0.05 [-0.13, 0.03] | **0.53 [0.36, 0.67]** | **0.23 [0.05, 0.42]** | 0.11 [-0.02, 0.29] | **0.37 [0.07, 0.62]** | -0.04 [-0.21, 0.12] | 0.02 [-0.10, 0.15] | 108 | 118 |
| *Strict (N=452)* | | | | | | | | | |
| *Phenotypic: Pearson correlation between GAD-7 and learning parameter (all individuals).* | | | | | | | | | |
| *Within-trait: twin correlation (Twin 1 vs Twin 2) for each measure separately.* | | | | | | | | | |
| *Cross-twin cross-trait: average of r(Twin1 GAD, Twin2 Param) and r(Twin1 Param, Twin2 GAD).* | | | | | | | | | |
| *95% CIs via percentile bootstrap (10000 resamples).* | | | | | | | | | |
| *Bold indicates 95% CI excludes zero.* | | | | | | | | | |

#### Supplementary table 12. Cross-twin cross-trait correlations (moderate exclusion criteria).

|  | **Phenotypic** | **Within-trait: GAD-7** | | **Within-trait: Parameter** | | **Cross-twin Cross-trait** | | **Pairs** | |
| --- | --- | --- | --- | --- | --- | --- | --- | --- | --- |
| **Learning Parameter** | **r [95% CI]** | **MZ r [95% CI]** | **DZ r [95% CI]** | **MZ r [95% CI]** | **DZ r [95% CI]** | **MZ r [95% CI]** | **DZ r [95% CI]** | **MZ n** | **DZ n** |
| Threat positive | **-0.10 [-0.18, -0.02]** | **0.49 [0.33, 0.62]** | **0.20 [0.03, 0.37]** | **0.18 [0.04, 0.31]** | 0.07 [-0.09, 0.22] | -0.03 [-0.16, 0.10] | -0.03 [-0.15, 0.09] | 138 | 160 |
| Threat negative | -0.03 [-0.11, 0.06] | **0.49 [0.33, 0.63]** | **0.20 [0.04, 0.36]** | 0.01 [-0.16, 0.23] | 0.11 [-0.10, 0.31] | 0.04 [-0.08, 0.17] | -0.03 [-0.13, 0.09] | 138 | 160 |
| Safety | -0.06 [-0.14, 0.02] | **0.49 [0.33, 0.62]** | **0.20 [0.03, 0.37]** | **0.24 [0.08, 0.40]** | 0.04 [-0.11, 0.21] | **-0.13 [-0.24, -0.03]** | -0.04 [-0.14, 0.06] | 138 | 160 |
| Threat extinction | **-0.18 [-0.26, -0.10]** | **0.49 [0.33, 0.63]** | **0.20 [0.04, 0.37]** | 0.14 [-0.01, 0.27] | 0.14 [-0.02, 0.29] | -0.07 [-0.19, 0.05] | -0.07 [-0.18, 0.03] | 138 | 160 |
| Safety confirmation | **-0.10 [-0.18, -0.01]** | **0.49 [0.33, 0.63]** | **0.20 [0.04, 0.36]** | **0.18 [0.03, 0.34]** | 0.06 [-0.09, 0.21] | -0.08 [-0.20, 0.04] | -0.02 [-0.13, 0.09] | 138 | 160 |
| CS+ Initial Value | 0.06 [-0.01, 0.13] | **0.49 [0.33, 0.63]** | **0.20 [0.04, 0.36]** | 0.00 [-0.16, 0.18] | 0.11 [-0.05, 0.27] | -0.10 [-0.21, 0.01] | 0.10 [-0.00, 0.20] | 138 | 160 |
| CS- Initial Value | 0.05 [-0.03, 0.13] | **0.49 [0.33, 0.63]** | **0.20 [0.04, 0.37]** | 0.04 [-0.13, 0.21] | 0.08 [-0.08, 0.25] | 0.04 [-0.08, 0.16] | 0.02 [-0.09, 0.13] | 138 | 160 |
| Extinction CS- Jump | -0.02 [-0.10, 0.05] | **0.49 [0.33, 0.62]** | **0.20 [0.03, 0.37]** | **0.28 [0.11, 0.44]** | 0.06 [-0.10, 0.22] | -0.03 [-0.15, 0.10] | 0.07 [-0.04, 0.18] | 138 | 160 |
| Response precision | -0.05 [-0.13, 0.01] | **0.49 [0.33, 0.63]** | **0.20 [0.04, 0.36]** | 0.11 [-0.00, 0.26] | **0.38 [0.15, 0.59]** | 0.00 [-0.14, 0.13] | -0.01 [-0.12, 0.10] | 138 | 160 |
| *Moderate (N=596)* | | | | | | | | | |
| *Phenotypic: Pearson correlation between GAD-7 and learning parameter (all individuals).* | | | | | | | | | |
| *Within-trait: twin correlation (Twin 1 vs Twin 2) for each measure separately.* | | | | | | | | | |
| *Cross-twin cross-trait: average of r(Twin1 GAD, Twin2 Param) and r(Twin1 Param, Twin2 GAD).* | | | | | | | | | |
| *95% CIs via percentile bootstrap (10000 resamples).* | | | | | | | | | |
| *Bold indicates 95% CI excludes zero.* | | | | | | | | | |

#### Supplementary table 13. Bivariate ACE model fitting.

| **Twin group** | **Model** | **-2LL** | **df** | **AIC** | **BIC** | **χ² vs ACE** | **p vs ACE** | **χ² vs AE** | **p vs AE** | **χ² vs CE** | **p vs CE** |
| --- | --- | --- | --- | --- | --- | --- | --- | --- | --- | --- | --- |
| Safety | | | | | | | | | | | |
| *Strict (N=452)* | ACE | 2512.12 | 893 | 726.12 | -2328.42 | - | - | - | - | - | - |
|  | **AE** | **2512.83** | **896** | **720.83** | **-2343.97** | **0.72** | **0.869** | **-** | **-** | **-** | **-** |
|  | CE | 2519.11 | 896 | 727.11 | -2337.68 | 7.00 | 0.072 | - | - | - | - |
|  | E | 2561.42 | 899 | 763.42 | -2311.64 | *49.31* | *<.001* | *48.59* | *<.001* | *42.31* | *<.001* |
| *Moderate (N=596)* | ACE | 3322.82 | 1181 | 960.82 | -3405.45 | - | - | - | - | - | - |
|  | **AE** | **3322.93** | **1184** | **954.93** | **-3422.43** | **0.11** | **0.991** | **-** | **-** | **-** | **-** |
|  | CE | 3331.66 | 1184 | 963.66 | -3413.70 | *8.84* | *0.031* | - | - | - | - |
|  | E | 3376.87 | 1187 | 1002.87 | -3385.58 | *54.05* | *<.001* | *53.94* | *<.001* | *45.21* | *<.001* |
| Threat extinction | | | | | | | | | | | |
| *Strict (N=452)* | ACE | 2505.97 | 893 | 719.97 | -2334.56 | - | - | - | - | - | - |
|  | **AE** | **2507.34** | **896** | **715.34** | **-2349.46** | **1.36** | **0.715** | **-** | **-** | **-** | **-** |
|  | CE | 2514.76 | 896 | 722.76 | -2342.04 | *8.78* | *0.032* | - | - | - | - |
|  | E | 2551.91 | 899 | 753.91 | -2321.15 | *45.94* | *<.001* | *44.58* | *<.001* | *37.16* | *<.001* |
| *Moderate (N=596)* | ACE | 3309.92 | 1181 | 947.92 | -3418.34 | - | - | - | - | - | - |
|  | **AE** | **3310.81** | **1184** | **942.81** | **-3434.54** | **0.89** | **0.828** | **-** | **-** | **-** | **-** |
|  | CE | 3317.96 | 1184 | 949.96 | -3427.40 | *8.04* | *0.045* | - | - | - | - |
|  | E | 3359.58 | 1187 | 985.58 | -3402.87 | *49.66* | *<.001* | *48.77* | *<.001* | *41.62* | *<.001* |
| *GAD-7 x Safety. Best-fitting model in bold (hierarchical LRT).Italic χ²/p indicates significantly worse fit (p < .05).* | | | | | | | | | | |  |

#### Supplementary table 14. Bivariate ACE model variance estimates.

|  | | | **GAD-7** | | **Parameter** | | **Cross-trait r** | |
| --- | --- | --- | --- | --- | --- | --- | --- | --- |
| **Learning Parameter** | **Twin Group** | **Model** | **A²** | **E²** | **A²** | **E²** | **rA** | **rE** |
| Safety | Strict (N=452) | AE | **0.509 (0.359, 0.649)** | **0.491 (0.351, 0.641)** | **0.155 (0.011, 0.304)** | **0.845 (0.696, 0.989)** | -0.242 (-1.049, 0.149) | 0.105 (-0.071, 0.258) |
|  | Moderate (N=596) | AE | **0.462 (0.322, 0.584)** | **0.538 (0.416, 0.678)** | **0.201 (0.067, 0.334)** | **0.799 (0.666, 0.933)** | -0.309 (-0.819, 0.007) | 0.062 (-0.097, 0.208) |
| Threat extinction | Strict (N=452) | AE | **0.516 (0.370, 0.640)** | **0.484 (0.360, 0.630)** | 0.104 (0.000, 0.251) | **0.896 (0.749, 1.000)** | -0.146 (-11294629.319, 1.260) | **-0.167 (-0.319, -0.017)** |
|  | Moderate (N=596) | AE | **0.466 (0.328, 0.604)** | **0.534 (0.396, 0.672)** | **0.179 (0.043, 0.307)** | **0.821 (0.693, 0.957)** | -0.275 (-0.686, 0.086) | **-0.148 (-0.268, -0.012)** |
| *Bivariate ACE twin models: GAD-7 x each learning parameter.* | | | | | | | | |
| *Best-fitting model shown (hierarchical LRT). Bootstrap 95% CIs in parentheses.* | | | | | | | | |
| *A² = additive genetic; E² = unique environment + error.* | | | | | | | | |
| *rA = genetic correlation; rE = unique-environment correlation.* | | | | | | | | |
| *Bold indicates 95% CI excludes zero.* | | | | | | | | |
